# A high-throughput sequencing approach identifies immunotherapeutic targets for bacterial meningitis in neonates

**DOI:** 10.1101/2022.12.22.521560

**Authors:** Stéphanie Pons, Eric Frapy, Youssouf Sereme, Charlotte Gaultier, François Lebreton, Andrea Kropec, Olga Danilchanka, Laura Schlemmer, Cécile Schrimpf, Margaux Allain, François Angoulvant, Hervé Lecuyer, Stéphane Bonacorsi, Hugues Aschard, Harry Sokol, Colette Cywes-Bentley, John J. Mekalanos, Thomas Guillard, Gerald B. Pier, Damien Roux, David Skurnik

**Author notes:** Corresponding author: Professor David Skurnik, Université de Paris- Hôpital Necker Enfants Malades, Département de Microbiologie, 149, rue de Sèvres, 75015 PARIS, FRANCE.

## Abstract

**Background:** Worldwide, *Escherichia coli* is the leading cause of neonatal Gram-negative bacterial meningitis, but full understanding of the pathogenesis of this disease is not yet achieved. Moreover, to date, no vaccine is available against bacterial neonatal meningitis.

**Methods:** Here, we used Transposon Sequencing of saturated banks of mutants (TnSeq) to evaluate *E. coli* K1 genetic fitness in murine neonatal meningitis. We identified *E. coli* K1 genes encoding for factors important for systemic dissemination and brain infection, and focused on products with a likely outer-membrane or extra-cellular localization, as these are potential vaccine candidates. We used *in vitro* and *in vivo* models to study the efficacy of active and passive immunization.

**Results:** We selected for further study the conserved surface polysaccharide Poly-β-(1-6)-N-Acetyl Glucosamine (PNAG), as a strong candidate for vaccine development. We found that PNAG was a virulence factor in our animal model. We showed that both passive and active immunization successfully prevented and/or treated meningitis caused by *E. coli* K1 in neonatal mice. We found an excellent opsonophagocytic killing activity of the antibodies to PNAG and *in vitro* these antibodies were also able to decrease binding, invasion and crossing of *E. coli* K1 through two blood brain barrier cell lines. Finally, to reinforce the potential of PNAG as a vaccine candidate in bacterial neonatal meningitis, we demonstrated that Group B *Streptococcus*, the main cause of neonatal meningitis in developed countries, also produced PNAG and that antibodies to PNAG could protect *in vitro* and *in vivo* against this major neonatal pathogen.

**Interpretation:** Altogether, these results indicate the utility of a high-throughput DNA sequencing method to identify potential immunotherapy targets for a pathogen, including in this study a potential broad-spectrum target for prevention of neonatal bacterial infections.

**Fundings:** ANR Seq-N-Vaq, Charles Hood Foundation, Hearst Foundation. Groupe Pasteur Mutualité

## Introduction

Neonatal bacterial meningitis continues to be a serious disease and even when treated with appropriate antibiotics, mortality rates over 10% still ensue. While Group B *Streptococcus* (GBS), remains the most common cause of neonatal sepsis and meningitis, worldwide, *Escherichia coli* strains causing neonatal meningitis (NMEC) are responsible for 15-20% of early-onset infections in term infants and for about 35% in preterm newborns.^1^ In addition, due to the emergence of antibiotic resistant *E. coli* K1 strains, notably those producing the extended spectrum beta-lactamase CTX-M, the mortality rates may further increase significantly in the future.^2^ Therefore, new modes of prevention and treatment are warranted based on an improved understanding of the disease pathophysiology and of *E. coli* K1 pathogenic factors. Using saturated transposon (Tn) mutagenesis and high-throughput sequencing (TnSeq), a powerful tool to investigate host-pathogen interactions, we fill-in-gaps in our knowledge about the contribution to overall fitness of the multitude of genes in *E. coli* K1 needed to cause neonatal meningitis. This approach led to the identification of PNAG as a strong candidate antigen produced by this pathogen that could be targeted by immunotherapy as PNAG was found to be both a virulence factor and to have an outer-membrane localization. We therefore fully explored this candidate, using both *in vitro* and *in vivo* studies.

## Methods

The detailed methods are available in supplementary materials.

## Results

### TnSeq and E. coli K1 essential genes

In this study, we initially selected the NMEC sequenced strain *E. coli* K1 S88, previously reported as an important cause of neonatal meningitis and belonging to the O-serotype 45.^3^ A bank of 3×10^5^ Tn-mutants of *E. coli* S88 was produced, grown overnight in lysogeny broth (LB) and the DNA recovered and subjected to high-throughput sequencing to identify sites and frequencies of Tn inserts into the genome (TnSeq). As previously reported with another pathogen, *Pseudomonas aeruginosa*,^4^ this allowed us to identify genes with no Tn inserts and thus considered as essential for the growth of *E. coli* K1 S88 in LB (Figure S1). To compare these results with other *E. coli* strains, a Tn-bank of 3×10^5^ mutants in the genome of the well described commensal strain *E. coli* K12 MG1655 and a Tn-bank of 3×10^5^ inserts in the *E. coli* LF82 strain,^5^ associated with inflammatory bowel disease, were also built and analyzed. This approach allowed us to have a better view of the essential genes in different strains of *E. coli* (Figure 1). Overall, we found 284 common essential genes in these three *E. coli* strains, close to the prevalence to the common essential genes found between two *P. aeruginosa* strains, PA01 and PA14^4^ (Table S1).

**Figure 1.**
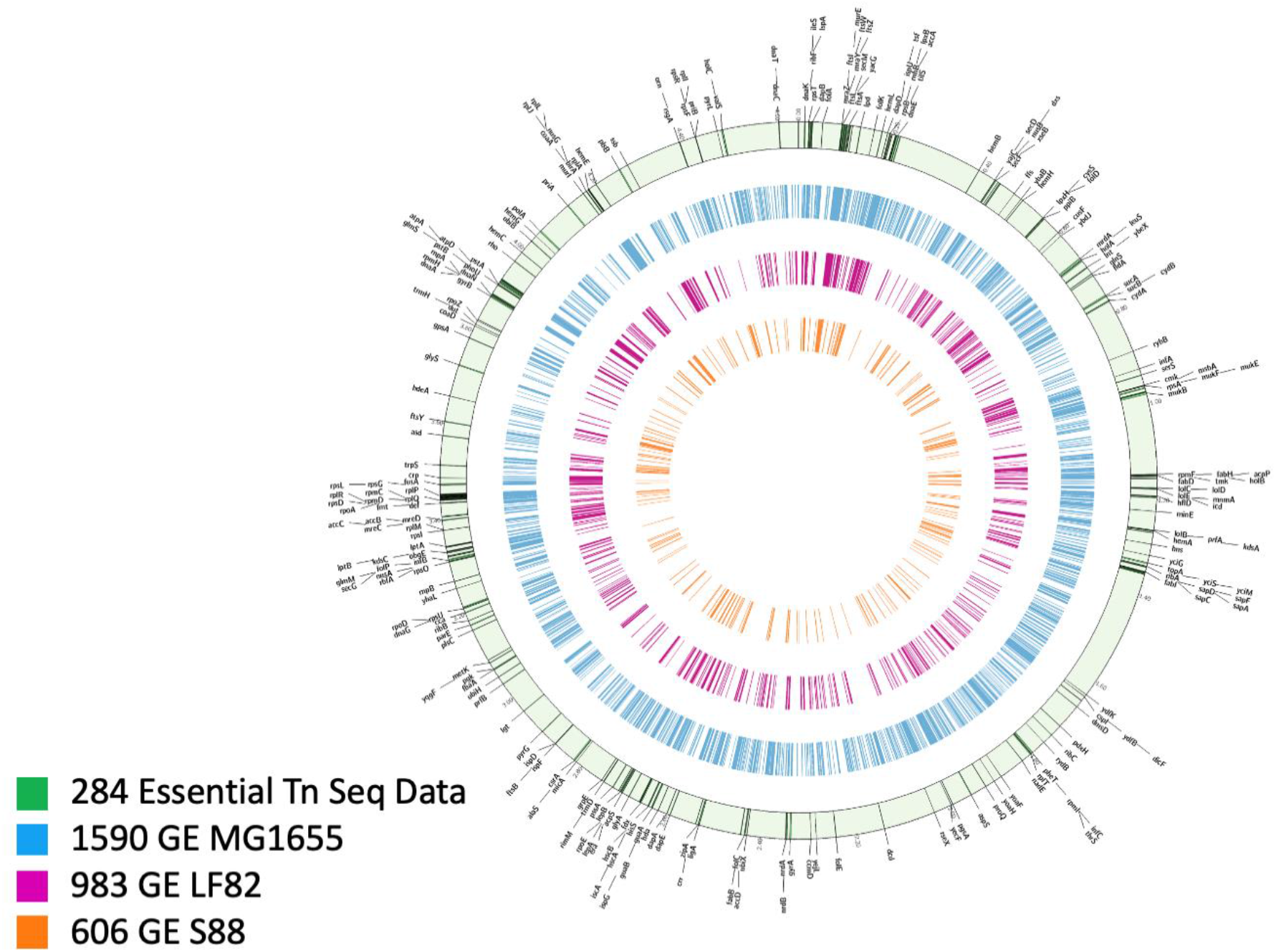
Comparative identification of essential genes in *E. coli*. After TnSeq analysis of three saturated bank of mutants of *E. coli* grown in LB, 1590 genes have been identified as essential in strain *E. coli* K12 MG1655 (purple circle), 983 in *E. coli* LF82 (blue circle) and 606 in *E. coli* K1 S88 (brown circle) for a total of 284 essential genes found in common for these three strains (annotated and located on *E. coli* MG1655 chromosome (green circle). Table S1 gives a complete listing of the essential genes found for these three strains in these conditions.

### *TnSeq using* E. coli *K1 in two mouse models of neonatal infection*

In order to investigate *E. coli* K1 pathophysiology, we established two mouse models of neonatal infection (Figure S2). The first *in vivo* model used 2 to 3 day-old mice given an intraperitoneal (IP) challenge with 5×10^6^ Colony Forming Units (CFU)/mouse of the Tn-bank that generated a high level of bacteremia. After 24h, the neonatal mice were sacrificed, and the livers, spleens, brains and meninges harvested to determine systemic dissemination followed by brain infection (above 10^8^ CFU/organ, Figure S3). The second model was designed to mimic the steps of pathogenesis of neonatal meningitis caused by *E. coli* K1. Neonatal mice were challenged by oral gavage, as the gastro-intestinal (GI) tract is likely the main site of initial colonization and origin for meningitis pathogens in neonates.^6^ After 24h, the neonatal mice were sacrificed and the livers, spleens, brains and meninges harvested (Figure S4). These two models allowed determinations of (i) genes important for systemic dissemination, following either IP infection or after GI tract colonization (both routes are seen in human infection), and (ii) genes important for brain infection following the route of infection thought to occur in the human disease (from the GI tract to the brain).

### Genes important for systemic dissemination using two routes of infection

To identify genes producing factors important for systemic dissemination of *E. coli* K1, the data generated from these TnSeq experiments were analyzed using the systematic approach we previously reported.^4,7^ Following the same approach here, analysis at the genome scale (Figure S5) revealed that selective pressures in the gavage model were greater than those in the IP model. This result was somewhat expected as the gavage model has an additional selective step compared to the IP model: the colonization of the GI tract. While it has been recently reported that this step could lead to anatomical bottlenecks, this anatomical barrier is avoided using the IP route of infection.^8^ Therefore, only mutants with a very strong decrease in fitness (at least a 10-fold decrease in the prevalence of the Tn-mutants between the *in vitro* and the *in vivo* conditions) using both routes of infection (gavage and IP) were further studied for follow-up analyses. Overall 179 genes in *E. coli* K1 S88 were considered important for systemic dissemination (Table S2). As shown (Figure 2, Figure S7 and Table S2), among the operons and individual genes having >10-fold changes in the output populations, genes encoding for major *E. coli* K1 virulence factors such as the K1 capsule^9^, the LPS, and OmpA^10^ were all confirmed to be important for systemic dissemination.

**Figure 2.**
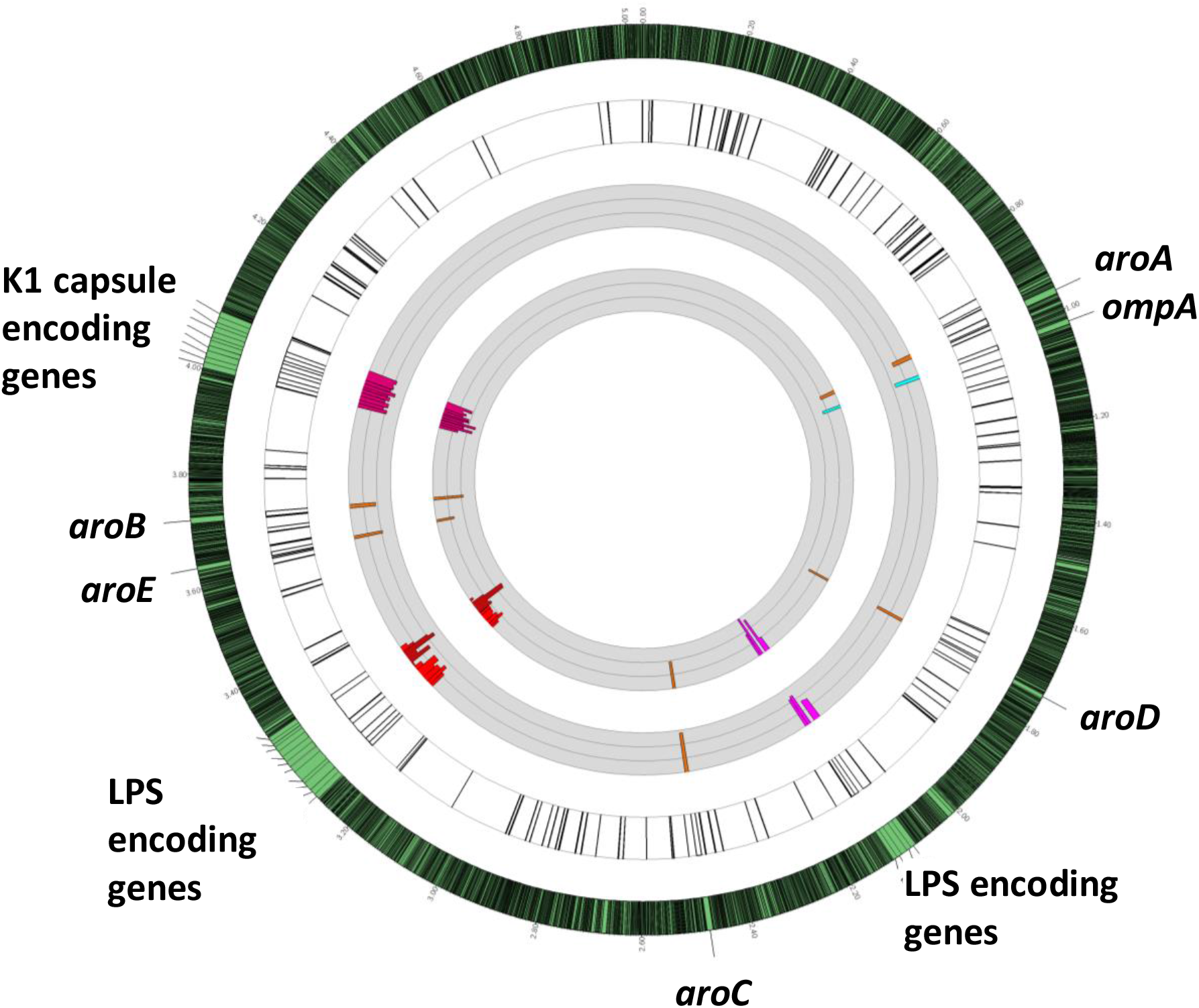
*E. coli* K1 *in vivo* (systemic dissemination) genetic fitness analysis using TnSeq localization on *E. coli* K1 S88 chromosome and quantitative analysis. **The outermost circle** represents the full *E. coli* K1 S88 genome with a 30-times magnification of the regions of interest. **The first white circle** represents the localization on *E. coli* K1 S88 chromosome of all the genes found important or essential for systemic dissemination (detailed Fig. S8 and Table S1) by TnSeq. Major, previously described, virulence factors of *E. coli* K1 are highlighted in color: LPS and K1 capsule encoding genes as examples for genes clusters, OmpA as example for a unique gene and Aro encoding genes as example of several individual genes (i.e not in cluster) located on several and distanced part of the chromosome. **The first gray circle** represents the fold change of the reads of the Tn insertions from the LB to the Liver after IP challenge. **The second gray circle represents** the fold change of the reads of the Tn insertions from the LB to the liver after gavage. **The thin grey circular lines** represent 10-fold-changes (i.e., at log_10_ scale). The limit of the fold change representation is 1,000. Bars represent changes in individual genes. Bars pointing toward the circle’s center represent Tn interrupted genes resulting in negative fitness.

While there is great interest in identifying *E. coli* K1 genes important for systemic dissemination,^8^ the main goal of this work was to study neonatal bacterial meningitis to identify vaccine candidate in this specific setting. Therefore, to identify the genes likely to be important or essential for brain infection in murine neonatal meningitis, we selected those with Tn-insertions leading to at least a 10-fold reduction in occurrence compared to *in vitro* growth in LB that also had a greater impact on brain infection compared to systemic dissemination.

### Genes important for brain infection

These genes were determined by being at least a 10-fold decreased in representation of the Tn-interrupted genes in the brain compared with the liver. The systematic analysis of the 1203 Tn-mutants with reduced fitness in brain infection under these conditions was undertaken (Table S3), focusing on those encoding for cell surface factors, outer membrane proteins or extracellular products (Figure 3). These are strong vaccine candidates due to the potential ability to raise protective antibodies to surface-accessible or extracellular antigens. As shown in Figure 3, Tn-inserts in four gene clusters corresponding to these criteria were found to have a >10-fold reduced occurrence in the brains of infected mice: the *fim* genes, the *pap* genes, the *csg* genes, and the *pga* genes. These encode for the type I Fimbriae, the P Fimbriae, Curli and the Poly-β-(1-6)-*N*-Acetyl-Glucosamine (PNAG) surface polysaccharide, respectively. Finding through our TnSeq experiments, and in our animal model, that PNAG encoding genes were among the few operons important or even essential for *E. coli* K1 neonatal meningitis pathophysiology was considered as an excellent and excellent opportunity. Indeed, this surface polysaccharide with capsule-like properties has previously been reported as a potential vaccine antigen in other infectious settings.^11^ Therefore the *pga* locus, its product, PNAG, and antibodies to PNAG were selected as best candidates to develop an efficient immunotherapy (both active and passive vaccination) against neonatal bacterial meningitis caused by *E. coli* K1.

**Figure 3.**
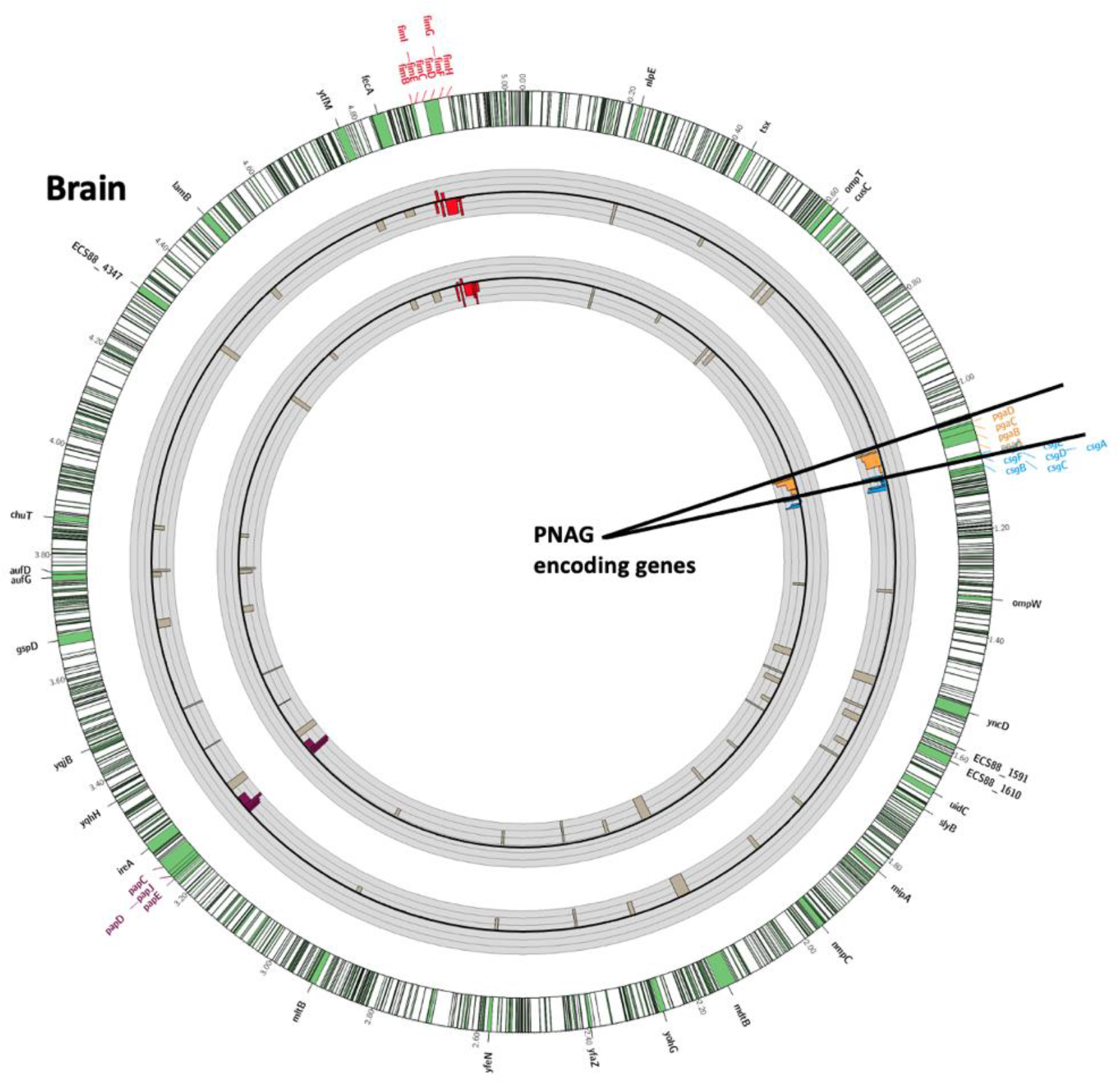
*In vivo* (brain infection) genetic fitness analysis using TnSeq of *E. coli* K1 mutants in genes encoding extracellular products or outer-membrane proteins. **The outermost circle** represents the full *E. coli* K1 S88 genome with a 30-times magnification of the regions of interest shown as green portions of the chromosome. **The first gray circle** represents the fold change of the reads of the Tn insertions from the Liver to the Brain after gavage. **The second gray circle** represents the fold change of the reads of the Tn insertions from the LB to the Brain after gavage. **Black circular lines in the middle of the grey circles** represent baseline. **Thin grey circular lines above and below the central black circular line** represent 10-fold-changes (i.e., a log_10_ scale). The limit of the fold change representation is 1,000. Bars represent changes in individual genes. Bars pointing toward the circle’s center represent Tn interrupted genes resulting in negative fitness whereas bars pointed outward represent genes whose loss increases fitness. Poly-β-(1-6)-N-Acetyl-Glucosamine (PNAG), Curly, Type 1 Fimbriae and Type 1 pilus encoding genes are highlighted in red, purple, blue and light brown respectively.

### *PNAG is a virulence factor of* E. coli *K1 and a potential vaccine target*

To confirm the data generated using TnSeq, and more precisely the role of PNAG as a fitness factor for *E. coli* K1 neonatal meningitis, we selected a strain belonging to the major NMEC O-serotype 18, *E. coli* K1 clinical isolate *E. coli* E11. We then deleted the *pga* genes in E11 (E11Δ*pga*) and used E11*Dpga* and E11 wild type for further analysis. Comparing bacterial burdens of wild-type and E11Δ*pga* recovered from the brains of infected neonatal mice (Figure 4A), significantly fewer *E. coli* E11Δ*pga* were found compared to the wild type E11 strain (*P*<0·01), highlighting the lower virulence of the Δ*pga* mutant and identifying PNAG as a virulence factor in our model. We next determined that PNAG was expressed on the surface of different strains of *E. coli* K1, including a multi-drug resistant strain producing the extended spectrum beta-lactamase (ESBL) CTX-M ^2^(Figure 4B and Figure S9A-D). PNAG production was detected on the surface of the NMEC *E. coli* K1 both when grown *in vitro* in LB and when recovered from *in vivo* infected brains of neonatal mice (Figure 4B-C), indicating that PNAG was a potential vaccine target for *E. coli* K1 strains. The use of an *E. coli* K11Δ*pga* KO strain (Figure 4B) confirmed the specificity of the polyclonal antibodies and the monoclonal antibody (MAb).

**Figure 4.**
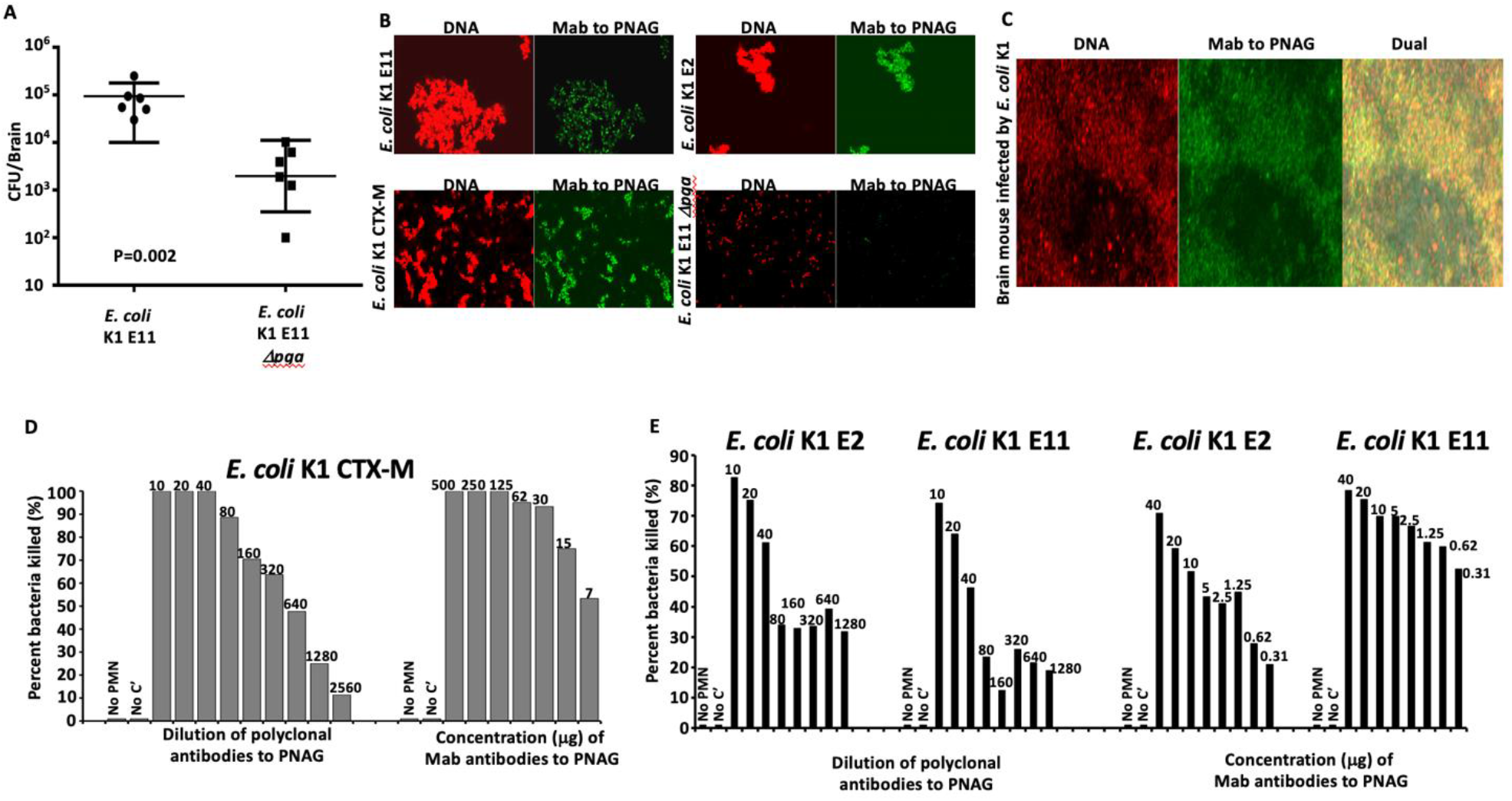
*E. coli* K1 and PNAG. **(A)** Loss of PNAG production (confirmed by confocal microscopy) in *E. coli* K1 (Δ*pga*) resulted in reduction from a median of 70,000 CFU/brain in wild—type (WT) to 3070 CFU/brain, P=0.002. Inoculum: 10^6^ CFU/mouse. **(B-C)** *In vitro* (**B**) and *in vivo* (**C**) detection of PNAG production (green) and bacterial DNA (red) by different strains of *E. coli* K1 using confocal microscopy to visualize binding of MAb F598 to PNAG. Bacteria in each microscopic field were visualized with a DNA stain, Syto83. **(D-E)** Percent killing of *E. coli* K1 producing the extended spectrum beta-lactamase CTX-M (**D**), *E. coli* K1 E2 and E11 (**E**) mediated by goat polyclonal antibodies to dPNAG-TT (anti-dPNAG) or a human MAb to PNAG (MAb F598). Killing calculated in comparison to CFU in corresponding tubes that contain normal goat serum or an irrelevant human IgG1 MAb. No bacterial killing is detected in the controls without leukocytes (No PMN) or without complement (No C’). Killing >30% is significant at P<0.05 using ANOVA and multiple-comparisons test.

### *In vitro activity of antibodies to PNAG against* E. coli *K1*

An opsonophagocytosis killing assay (OPK) was used to test the functional activity of antibodies to PNAG against *E. coli* K1, using both polyclonal antibodies^12^ and a fully human IgG1 MAb^13^. As shown in Figure 4D-E, both polyclonal antibodies and the MAb to PNAG were able to kill different strains of NMEC in the presence of complement and phagocytes in a dose-dependent manner.

### *Prevention of neonatal meningitis caused by* E. coli *K1 provided by antibodies to PNAG*

The prophylactic activity of PNAG antibodies was tested in the mouse model of neonatal meningitis, administering the challenge by oral gavage using four different strains of *E. coli* K1: S88, E2, E11 and CTX-M (Figure 5 and Figure S9E)^2^. Significant reduction of the bacterial load was found also in organs other than the brain (Figure S10). Both polyclonal antibodies and the human MAb to PNAG injected 24h before infection resulted in a significant decrease in the CFU recovered from the mice infected by S88, E2 and E11 (*P*<0·01, *P*<0·01, and *P*<0·01 respectively). When using the *E. coli* K1 strain producing the ESBL CTX-M, we found this organism led to a moribund/lethal infection in less than 24 hours in non-protected controls, but nonetheless injection of either polyclonal antibodies or the human MAb to PNAG resulted in significantly less neonatal deaths in 24 hours (*P*=0·01 and *P*<0·01, respectively).

**Figure 5.**
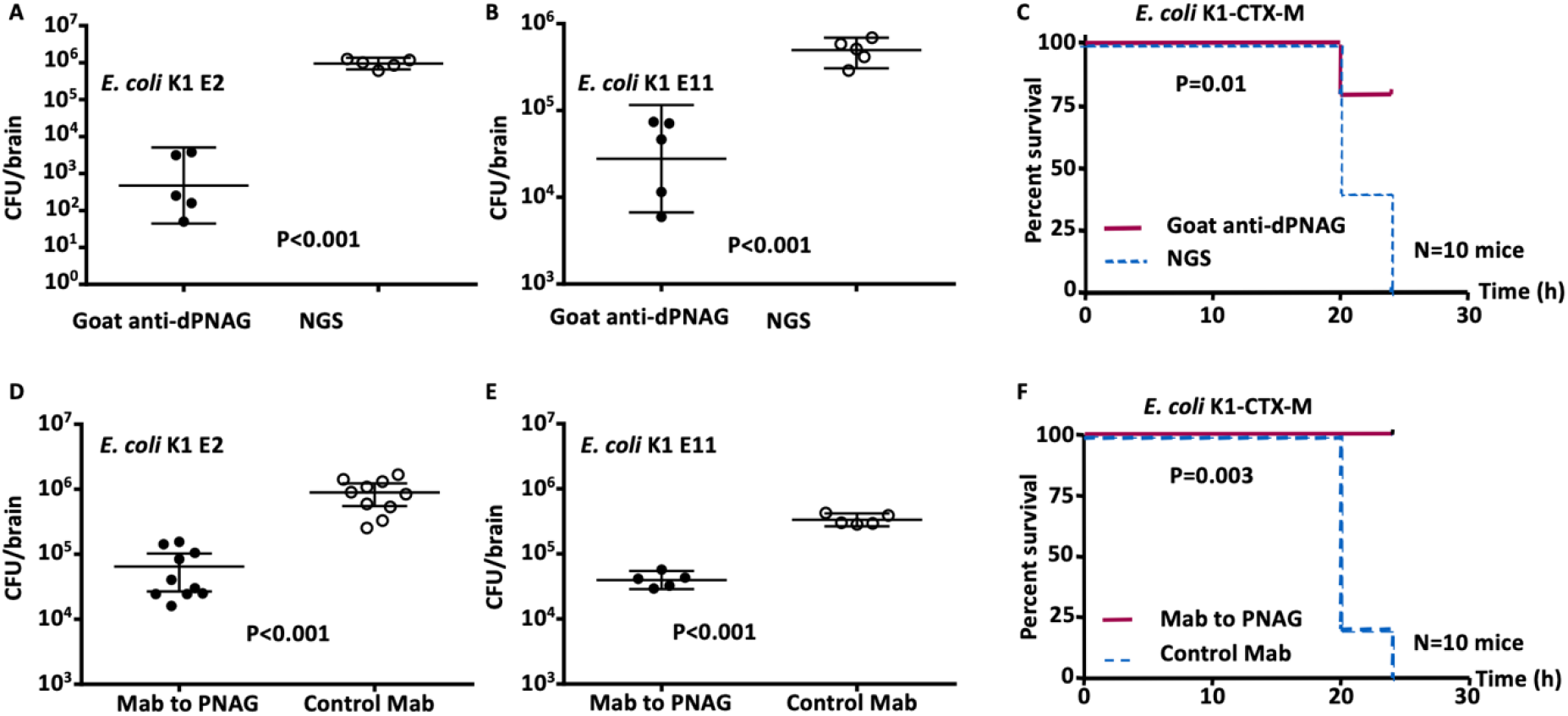
*In vivo* activity of antibodies to PNAG to protect against *E. coli* K1. Prophylactic (24 h pre-challenge) effect of 50 µl of opsonic goat polyclonal antibodies to PNAG (Goat anti-dPNAG) **(A-C)** or 50 µg of human MAb F598 to PNAG **(D-F)** on *E. coli* K1 levels in the brain (**A, B, D, E**) or on lethality rate (**C, F**) 24 h after challenge by gavage. Controls received normal goat serum (NGS) or an irrelevant human IgG1 MAb F429. Bars indicate medians. Challenge doses: 10^6^ CFU/animal (**A-B**), 10^5^ CFU/animal (**C-E**) and 10^4^ CFU/animal (**F**). P values determined by nonparametric t-test (**A, B, D, E**) and LogRank test (**C and F**). Each circle (**A, B, D, E**) represents one animal.

### *Treatment of severe neonatal infections caused by* E. coli *K1, with or without a co-infection caused by* Staphylococcus aureus

To try to mimic a clinical scenario in human neonatal meningitis diagnosis and treatment, we next tested the efficacy of antibody to PNAG to impact infection three hours subsequent to *E. coli* K1 infection. However, the causative pathogen in a human would take some time to be identified after the onset of signs of infection (whether it is indeed *E. coli* K1 or another pathogen). This could damper the potential use of PNAG antibodies after the start of the neonatal infections. However, several other PNAG producing pathogens, including potentially multi-drug resistant (MDR) bacteria such as *Staphylococcus epidermidis* and more particularly *S. aureus*, can also cause severe neonatal infections, in particular in premature neonates. ^14^ Therefore, to look for the therapeutic efficacy of antibodies to PNAG in a setting where there could be several different PNAG-producing MDR pathogens causing neonatal meningitis, neonatal mice were treated 3h after bacterial challenge without knowing if the animals were infected by *E. coli* K1 (gavage meningitis model), or by *S. aureus* (bacteremia induced by an intra-peritoneal challenge), or co-infected by both (unlikely clinical event, but tested as proof of concept for polymicrobial infection). We used several strains of *E. coli* K1 and *S. aureus* in this challenge setting, including the MDR CTX-M producing *E. coli* K1, and the methicillin-resistant *S. aureus* strain USA300 LAC.

As shown in Table 1 and Table 2, there was excellent therapeutic efficacy of both polyclonal antibodies and the human MAb to PNAG in reducing mortality and bacterial burdens regardless if the neonatal mice were infected by *E. coli* alone (*E. coli* K1 E11 or *E. coli* K1 CTX-M), *S. aureus* alone (*S. aureus* PS80 or *S. aureus* LAC), or co-infected by *E. coli* K1 and *S. aureus*. In some challenges a moribund/mortality outcome was used when the individual strain or combination caused this state to occur in <24 h, but in other challenges when neonates were still alive 24h after challenge (time point mandated for ending the experiment), CFU from the brain and the liver were counted after euthanasia. In both outcome scenarios, the antibodies to PNAG provided significant protection (*P<*0·01) (Table 2).

**Table 1.** Therapeutic activity of polyclonal and monoclonal antibodies to PNAG 3 hours after challenge of newborn mice with different *E. coli* K1 strains with and without *S. aureus* co-infection.

| Strains tested | Survivors/total challenged |  |
| --- | --- | --- |
|  | Goat polyclonal antibodies to PNAG | Normal Goat Serum |
| *E. coli* CTX-M | 5/5 | 0/5 |
| *E. coli* CTX-M + *S. aureus* PS80 | 5/5 | 0/5 |
| *E. coli* CTX-M + *S. aureus* LAC | 5/5 | 0/5 |
| *E. coli* E11 | 5/5 | 5/5* |
| *E. coli* E11 + *S. aureus* PS80 | 5/5 | 4/5* |
| *E. coli* E11 + *S. aureus* LAC | 4/5 | 0/5 |
|  | Monoclonal antibody to PNAG (F598) | Control monoclonal antibody (F429) |
| *E. coli* CTX-M | 4/5 | 0/5 |
| *E. coli* CTX-M + *S. aureus* PS80 | 5/10 | 0/10 |
| *E. coli* CTX-M + *S. aureus* LAC | 6/10 | 0/10 |
| *E. coli* E11 | 6/10 | 0/10 |
| *E. coli* E11 + *S. aureus* PS80 | 6/10 | 0/10 |
| *E. coli* E11 + *S. aureus* LAC | 6/10 | 0/10 |

\*Significant decrease of CFU in the brains

**Table 2.** Therapeutic activity of polyclonal and monoclonal antibodies to PNAG 1 hour after challenge of baby mice with two different *S. aureus* strains.

| Strains tested | Survivors/total challenged |  |
| --- | --- | --- |
|  | Goat polyclonal antibodies to PNAG | Normal Goat Serum |
| *S. aureus* LAC | 5/5 | 0/5 |
| *S. aureus* PS80 | 5/5 | 5/5* |
|  | Monoclonal antibody to PNAG (F598) | Control monoclonal antibody (F429) |
| *S. aureus* LAC | 5/10 | 0/10 |
| *S. aureus* PS80 | 6/10 | 0/10 |

\*Significant decrease of CFU in the livers

### Protection of neonates provided by transfer of maternal antibodies to PNAG

Another approach to prevent neonatal infections is *via* active vaccination of females, either prior to pregnancy, or in the case of humans, in the third trimester of pregnancy. To assess this means of protection, adult female mice were immunized either with a PNAG vaccine or a control antigen, the capsular polysaccharide 5 (CP5) of *S. aureus*.^15^ After immunization with PNAG, specific antibodies were successfully detected in the sera of vaccinated mice by ELISA (Figure 6). Then, mice born either from dams immunized with PNAG or CP5 were infected by gavage with *E. coli* K1. As shown in Figure 6B-C, significantly less bacteria were recovered from mice born to PNAG immunized dams (*P<*0·01). Next, we conducted a cross fostering experiment, placing newborn mice from dams immunized with CP5 in a cage with another dam immunized with CP5 or in a cage with a dam immunized with PNAG. Significant protection was observed in neonatal mice nursed by the PNAG immunized mother (*P*=0·02) compared to the neonatal mice nursed by a CP immunized dam, suggesting that transmission of antibodies to PNAG through the milk could be another source of protective antibodies in this model (Figure 6D-E). In this regard, it has recently been shown that IgG antibodies in milk can be transferred to the bloodstream of feeding pups *via* the neonatal Fc receptor.^16^

**Figure 6.**
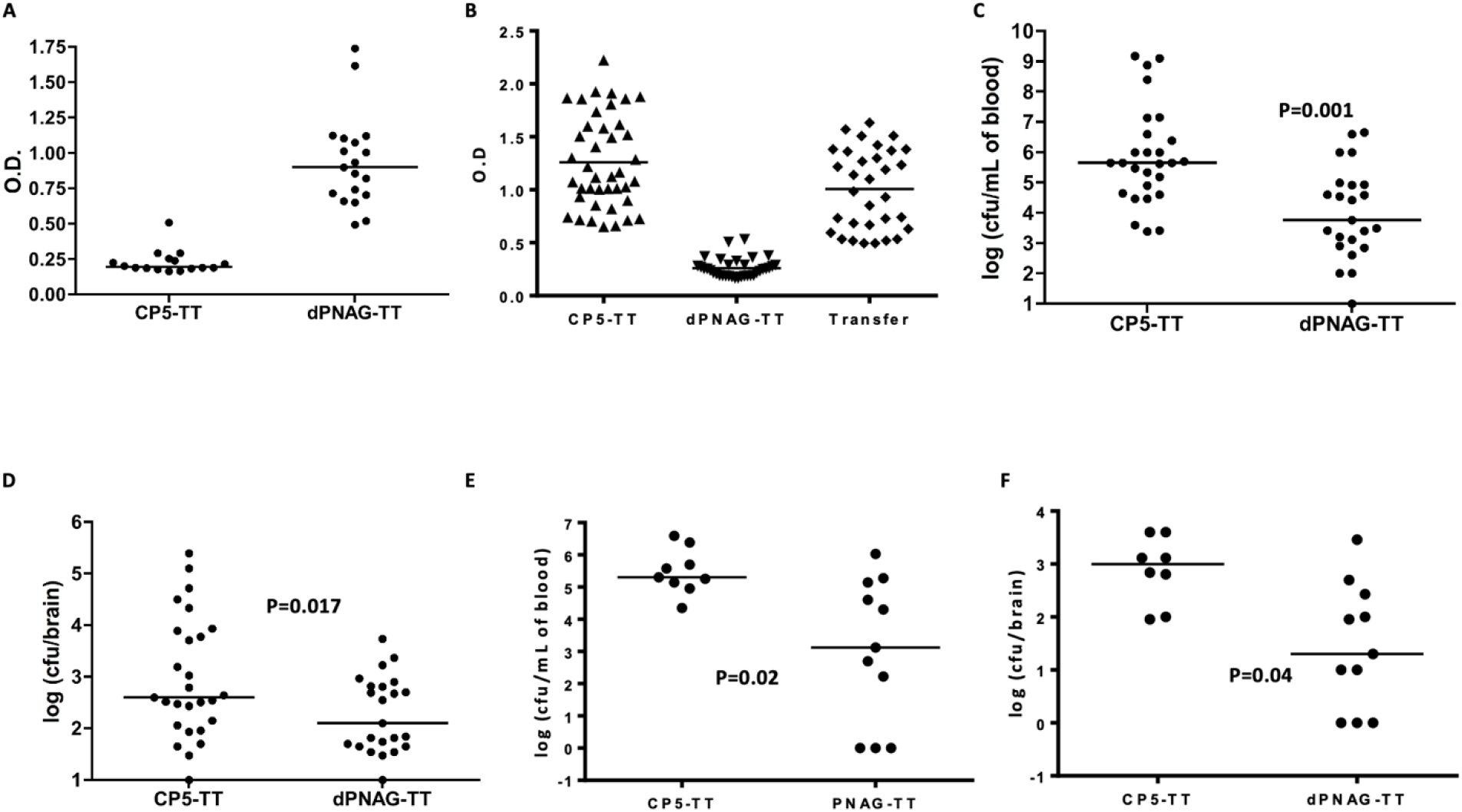
Protection provided by maternal antibodies to PNAG. Pups born from CP5-TT or dPNAG-TT immunized C3H/HeN female and infected with *E. coli* K1 S88 (10^6^ CFU/animal) by gavage. (**A-B)** Antibodies to PNAG were detected by ELISA in the serum of the mothers immunized by dPNAG-TT **(A)**, in the pups born from mothers immunized with dPNA-TTG or transferred in the cage of a mother immunized with dPNAG-TT **(B)**. Points=individual mice; lines= medians. (**C-D)** At 24h, significantly less CFU were recovered from the blood (**(C)** P=0.001, unpaired t-test) taken from the heart or from the brains (**(D)** P=0.017, unpaired t-test) of the neonates born from dPNAG-TT immunized mothers. Bars represent the median. Each circle represents an animal. **(E-F)** Pups born from CP5-TT immunized mother were placed at birth either in cages with dPNAG-TT immunized mothers or CP5-TT immunized mothers. Twenty-four hours after infection, significantly less CFU were recovered from the blood (**(E)** P=0,02, unpaired t-test) or from the brains (**(F)** P=0,04, unpaired t-test) of the neonates transferred in a cage with a mother immunized with dPNAG-TT. Bars represent the mean. Each circle represents an animal.

### *Antibodies to PNAG and crossing of the blood brain barrier (BBB) by* E. coli *K1*

To understand if mechanisms of antibody action other than OPK could also be active in protecting neonatal mice from meningitis, complement-C3 deficient neonatal mice were challenged in the neonatal meningitis murine protocol so that opsonic killing of *E. coli* by antibody would not be effective. C3-deficient neonatal mice were treated with polyclonal antibodies to PNAG then infected with *E. coli* K1 *via* gavage. No differences in the CFU levels were found in the livers and spleens (Figure 7A) of the C3-deficient mice immunized with antibodies to PNAG compared with those receiving the control antibodies. This result indicates that the OPK activity protects against *E. coli* K1 systemic dissemination. In contrast, there was a significant reduction in the CFU recovered from the brains of the C3-deficient mice injected with antibodies to PNAG compared to controls (*P<*0·01), suggesting an ability of these antibodies to block translocation of *E. coli* across the BBB.

**Figure 7.**
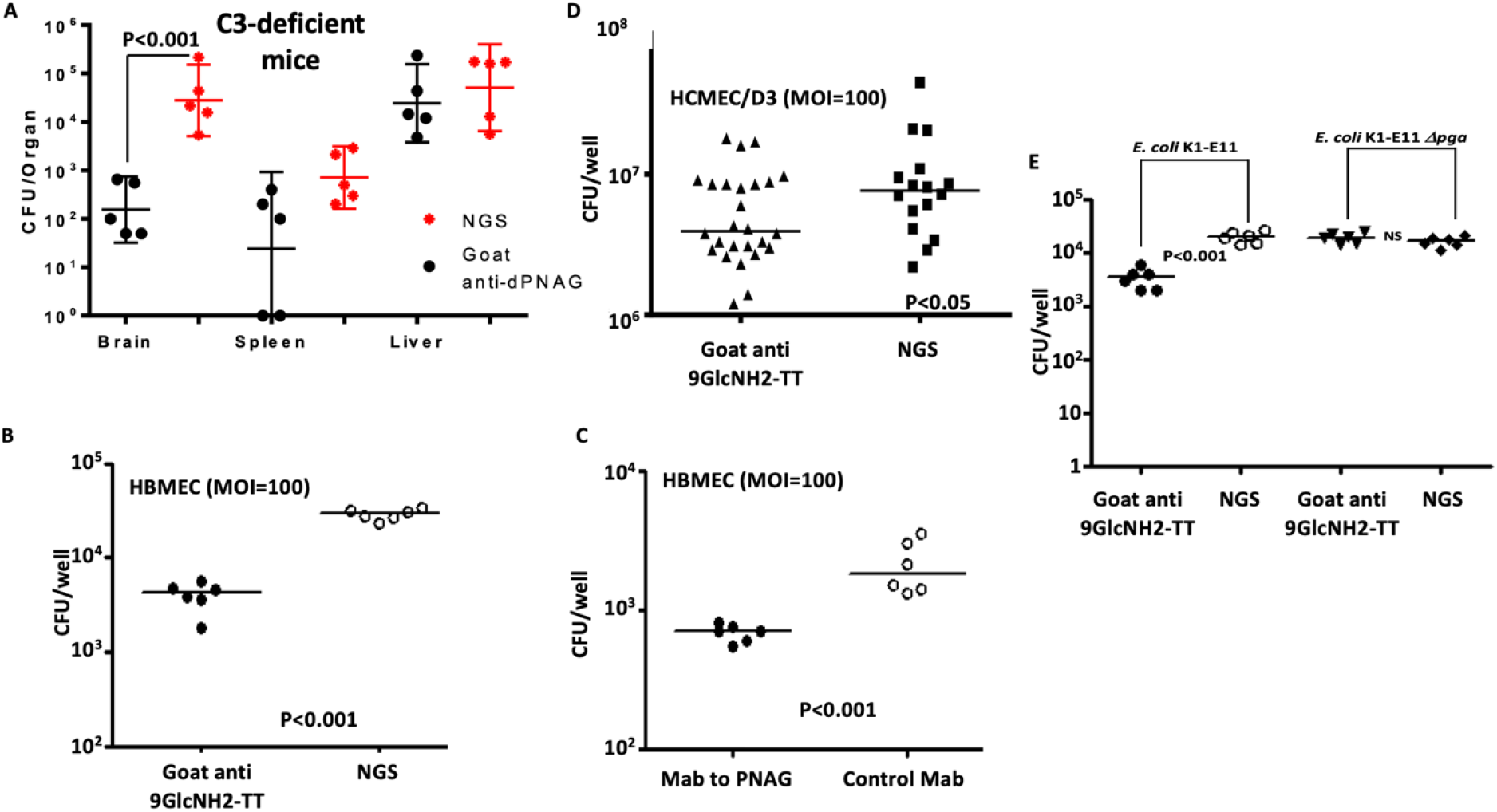
Antibodies to PNAG and crossing of the blood brain barrier. **(A)** Antibodies to PNAG significantly reduced bacterial levels in the brain (P<0.001) but not liver or spleen in C3-deficient mice. Points=individual mice; lines= medians. P value determined by non-parametric T-test. NGS=Normal goat serum. **(B-C)** Goat polyclonal antibodies **(B)** and human monoclonal antibodies **(C)** to PNAG inhibits adherence of *E. coli* K1 applied to HBMEC cultures. **(D)** Goat polyclonal antibodies to the synthetic oligosaccharide 9GlcNH2-TT significantly inhibits adherence of *E. coli* K1 E11 applied to HCMEC/D3 cultures. **(E)** No effect on adherence was obtained when using a Δ*pga E. coli* K1 E11 strain. Symbols are individual wells, lines medians, P values unpaired t-tests.

Next, we evaluated if this reduction in bacterial burdens in the brains of the C3-deficient mice could be linked to the inhibition of the binding, adhesion or translocation of *E. coli* K1 through the BBB conferred by the antibodies to PNAG. Two human cell lines were used to test this hypothesis: the Human Brain Micro Endothelial Cells (HBMEC) developed by KS Kim and the Human Cerebral Microvascular Endothelial Cells (HCMEC/D3) developed by PO Couraud.^17,18^ As shown in Figure 7B-D, regardless of the cell line used in the experiments, there was a significant reduction (*P<*0·01 and *P<*0·05, respectively) of *E. coli* K1 adhesion to the confluent monolayer cells in the presence of antibodies to PNAG. This result was obtained with polyclonal antibodies and the human MAb and with different strains of *E. coli* K1 (Figure S11). In addition, a significant reduction of invasion into cells in a monolayer and translocation through cells growing on transwell inserts was also found in the presence of antibodies to PNAG (Figure S11). However, this decreased translocation of *E. coli* through the BBB in presence of anti-PNAG antibodies was not secondary to a change in the monolayer integrity analyzed by trans-endothelial electrical resistance, as shown in Figure S12. To study the specificity of the PNAG antibodies in these *in vitro* experiments, we performed an assay using the parental *E. coli* K1 strain and the *E coli* K1 Δ*pga* isogenic mutant. No change of the adhesion to the HBMEC was observed with *E. coli* K1 Δ*pga* in the presence of antibodies to PNAG, establishing the specificity of the anti-PNAG activity. Taken together, these results suggest that in addition to their OPK activity, antibodies to PNAG may also protect against meningitis caused by *E. coli* K1 by interfering with the crossing of the BBB.

### *PNAG antibodies and Group B* Streptococci

Having found that PNAG antibodies were efficient in our models to treat infections caused by *E. coli* K1 and other PNAG producing pathogens, including MDR such as MRSA strains, we decided to look at the major cause of bacterial neonatal meningitis: *Streptococcus agalactiae* or Group B *streptococcus* (GBS).^19^ Indeed, if PNAG could also target GBS, it would probably significantly increase its potential clinical impact. In this context, first we successfully detected PNAG at the surface of GBS using confocal microscopy and the MAb to PNAG F598 and confirmed the properties of the binding antigen following the approach described by Cywes *et al*.^20^ by showing that the MAb binding remained after digestion with chitinase but was lost following digestion with the PNAG-degrading enzyme, Dispersin B, as well as following PNAG-hydrolyzing treatment with sodium periodate (Figure 8A-C).

**Figure 8.**
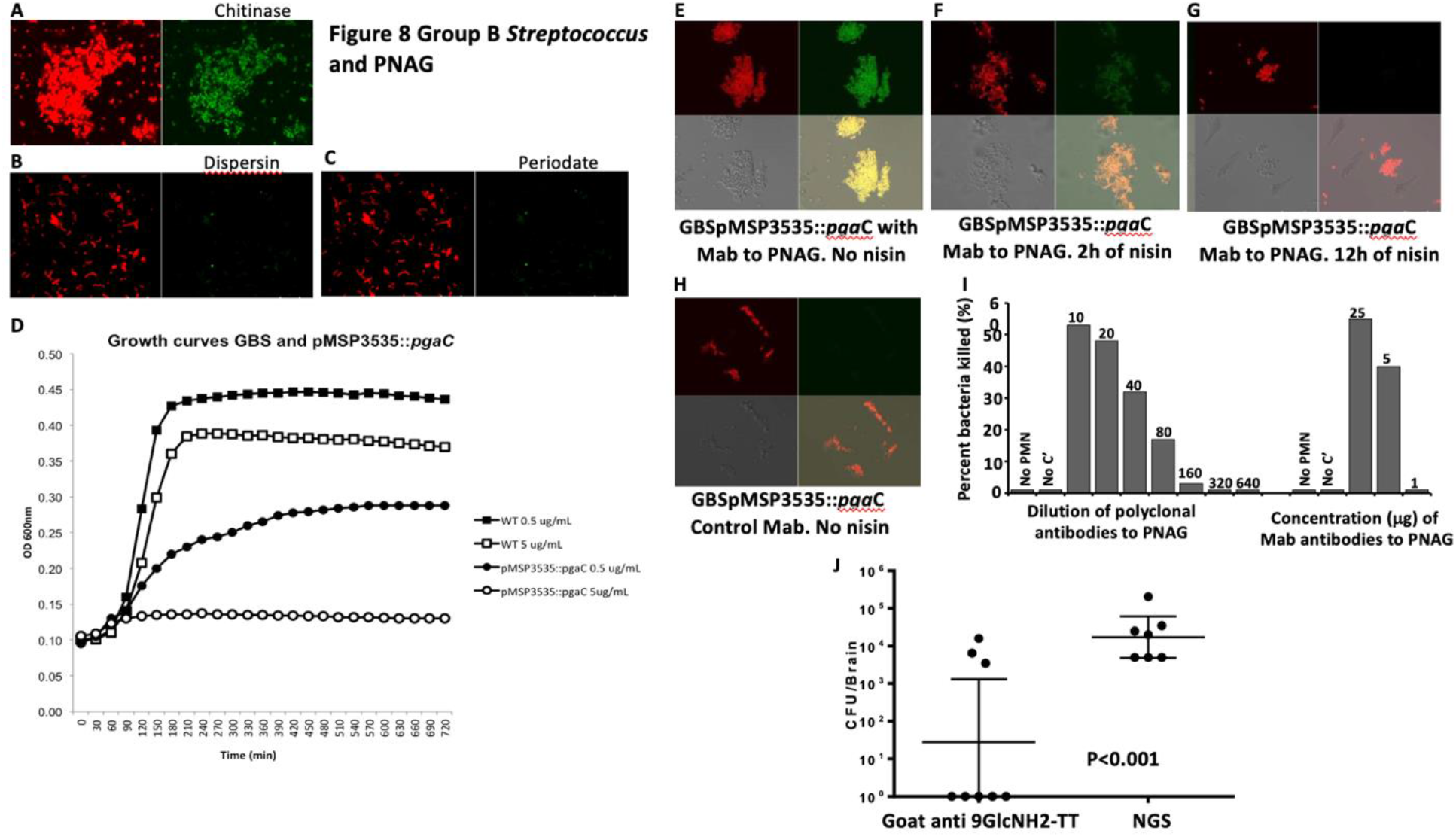
Group B streptococcus (GBS) and PNAG. **(A-C)** Detection of PNAG production (green) and bacterial DNA (red) by GBS using confocal microscopy to visualize binding of MAb F598 to PNAG. The binding is resistant to chitinase (**A**), susceptible to dispersin B (**B**) and to periodate (**C**). **(D)** GBS NEM 16 pMSP3535::*pga*C exhibits a decreased growth in presence of low concentration of nisin compared to the GBS NEM 16 (WT) cultured with low or high concentrations of nisin. A high concentration of nisin completely inhibits GBS pMSP3535::*pga*C growth. **(E-H)** Detection of PNAG production by GBS pMSP3535::pgaC grown in the absence (**E**) or the presence of nisin (**F**: 2h of nisin, **G**: 12h) using confocal microscopy. Green color: detection of the binding to PNAG by the Mab to PNAG (F598). Red color: bacterial DNA. The MAb F429 to *P. aeruginosa* alginate was used as control (**H**). **(I)** Killing of GBS by the polyclonal goat antiserum raised to a a nonameric β-1-6-linked glucosamine oligosaccharide conjugated to tetanus toxoid (9GlcNH_2_-TT) and of the fully human IgG1 MAb to PNAG, F598. NGS or MAb F429 to *P. aeruginosa* alginate were used as controls. **(J)** Goat polyclonal antibodies to the synthetic oligosaccharide 9GlcNH2-TT significantly decrease GBS CFU in the brains compared to NGS in a murine neonatal meningitis model after gavage. No CFU were detected in 5/8 of the neonates immunized with the PNAG antibodies. P value: unpaired t-test.

Next, a gene potentially involved in PNAG synthesis was identified in the sequenced GBS strain NEM316, encoding for a protein homologous to *E. coli pgaC* and designated GBS1605 (Blast *P* value=10e-24). Multiple attempts to truncate or delete GBS1605 by double cross overs using a temperature-sensitive shuttle vector^21^ were unsuccessful, suggesting that GBS1605 was an essential gene for GBS NEM316. Therefore, we used an antisense RNA (asRNA) approach to decrease the expression of GBS1605. A 721 bp asRNA, both complementary and overlapping with the ATG start codon of GBS1605, was cloned under the control of a nisin inducible promoter in the expression vector pMSP3535^22^. By qRT-PCR we found that increasing concentrations of nisin caused an increased expression of the asRNA and a corresponding decreased expression of GBS1605 transcripts (Figure S13). Concomitant with decreased GBS1605 mRNA expression, we observed that the growth of strain NEM316 was reduced (0·5 µg/ml nisin induction) or abolished (5 µg/ml nisin) (Figure 8D). Testing of PNAG production by GBS NEM316 WT carrying either the asRNA for GBS1605 or an empty vector control showed that decreased gene expression led to loss of PNAG production (Figure 8E-H). While the reliance of GBS NEM316 on a gene for both growth and PNAG production precluded testing the mutant for changes in virulence, this finding did encourage testing of the efficacy of antibodies to PNAG against GBS for opsonic killing activity and protective efficacy in the neonatal meningitis model. We found that antibodies to PNAG had potent OPK activity against GBS (Figure 8I). Additionally, significantly less GBS bacteria were recovered from the brains of neonatal mice passively immunized with antibodies to PNAG compared with those given control sera (*P<*0·01) (Figure 8J).

Thus, functional antibodies to PNAG have protective potential against the two major causes of neonatal bacterial meningitis (*E. coli* K1 and GBS) and less frequent cause of infections but more often associated with MDR such as Methicillin resistant *S. aureus*.

## Discussion

While antibiotic prescription is endlessly raising even during a viral pandemic such as the COVID-19 crisis^23^ that started in December 2019, new approaches to treat bacterial infections are urgently needed. In this context, the use of passive^24^ or active^25^ immunotherapies against microbial infections are more relevant now than ever. In this work, we were able to show that high-Throughput sequencing approaches can be used to identify targets and accelerate the process of vaccine development for bacterial infections. Prior strategies applied to the development of an effective vaccine against extra-intestinal pathogenic *E. coli* infections have not yet resulted in an effective vaccine for human use. This is partly due to the high antigenic heterogeneity of the major surface polysaccharides, LPS O and capsular K-antigens, that can be targeted for vaccination, making it very challenging to design a vaccine when targeting these carbohydrate.^26^ Outer membrane protein A was identified as important for systemic dissemination in the mouse model used here (Fig S6). While not selected for further investigation, OmpA has potential as a vaccine for neonatal meningitis.

In this work, we demonstrated that PNAG was a virulence factor for optimal neonatal meningitis caused by *E. coli* K1 and that antibodies to PNAG were able to prevent and treat *E. coli* K1 neonatal meningitis in a mouse model. In addition to their opsonophagocytic activity, these antibodies directly blocked the adhesion and translocation of *E. coli* across the BBB. A non-specific ability to prevent binding of a bacteria opsonized by antibodies targeting a surface polysaccharide has already been described. Indeed, this mechanism of action has been suggested as responsible for a decreased carriage of *S. pneumoniae* in patients vaccinated with Prevnar.^27^ We further found that the most frequent bacterial cause of neonatal meningitis, GBS, also produced PNAG and we were able to show protection against GBS, as well as a more uncommon cause of neonatal meningitis but often associated with antibiotic resistance, *S. aureus*. Two points seems of importance for the further development of antibodies to PNAG.

First, an initial conjugate vaccine for PNAG has been tested in humans for safety and immunogenicity^28^ and elicits broadly opsonic and bactericidal antibodies against a variety of pathogens.^29^ Second, the availability of a fully human IgG1 MAb that has also been in phase 1 and 2 human trials^30,31^ and is a potential immunotherapeutic for preventing neonatal meningitis, particularly in infants at high risk for infections. While PNAG is expressed by several commensal bacteria, no impact on the microbiota has been reported.^32^

These findings support further investigations of vaccine and MAb safety in pregnancy and neonates, which, if successful, could lead to testing of this potential means to prevent serious neurologic infections in neonates.

## Contributors

SP and DR performed *in vitro* and *in vivo* experiments, analyzed data and worked on the manuscript. EF analyzed data and edited the manuscript, CG performed *in vivo* experiments on *E. coli* K1 and edited the manuscript, FL worked on the genetics of GBS and edited the manuscript, AK performed *in vivo* experiments on *E. coli* K1 and edited the manuscript, OD performed high-throughput sequencing experiments and edited the manuscript, LS, SC, AM, AF LH, BS, HA et HS analyzed the data, CCB performed *in vitro* experiments on PNAG, JJM analyzed data and edited the manuscript, TG performed in vitro experiments on *E. coli* K1, analyzed the data and edited the manuscript, GBP analyzed the data, designed experiments, worked on the manuscript, D.S. developed the study concept, supervised the project, designed and performed *in vitro* and *in vivo* experiments on GBS, *S. aureus* and *E. coli* K1, analyzed data and wrote the manuscript.

## Data sharing

All data are available in the main text or the supplementary materials. All RPKM for 1162 irrelevant genes are available from the corresponding author upon reasonable request.

## Fundings

SP received an award from the Groupe Pasteur Mutualité

DR received an award from the Hearst Foundation

OD was supported by AI-026289 from National Institute of Allergy and Infectious Disease

CCB received an unrestricted gift from Alopexx Vaccines

DS received an award from the ANR (AAPG2020, France)

DS received an award from the Charles H. Hood Foundation

## Acknowledgments

None

## Supplementary Materials

Figures S1 to S13

Tables S1 to S5

## Supplementary materials

### Methods

#### Experimental design

##### Ethics statement

The Harvard Medical School and Brigham and Women’s Hospital animal management programs are accredited by the Association for the Assessment and Accreditation of Laboratory Animal Care, International (AAALAC), and meets National Institute of Health standards as set forth in the 8^th^ edition of the Guide for the Care and Use of Laboratory Animals (National Research Council “Animal Care and Use Program”. Guide for the Care and Use of Laboratory Animals: Eighth Edition. Washington, DC: The National Academies Press, 2011). The institution also accepts as mandatory the PHS Policy on Humane Care and Use of Laboratory Animals by Awardee Institutions and NIH Principles for the Utilization and Care of Vertebrate Animals used in Testing, Research, and Training. All animal studies conducted in this research were approved by the Institutional Animal Care and Use.

##### Strains

Strains used in this study are listed in Table S4. Primers and plasmids are listed Table S5.

##### E. coli *TnSeq*

Illumina libraries were prepared as described. Briefly, bacterial DNA was extracted, digested with MmeI, and ligated to oligonucleotide adapters, PCR amplified and sequenced using Illumina HiSeq.^1,2^ Between 20 and 50 million sequencing reads per sample were recovered. All sequences retrieved from the Illumina reactions were trimmed to eliminate those reads with quality scores less than 0·05 and/or sequences with ambiguous nucleotides.^1,2^ Data were analyzed using the RNA-seq module of the Bioinformatics Workbench software package (CLC) (http://www.clcbio.com/index.php?id=1240). Circos figures in the main text (Figure 1 and Figure 2) and in the supporting information figures (Figure S7) were drawn following the instructions provided at http://www.circos.ca/. For each Tn-insert, the significance of the difference in the proportions of the RPKM from one environment to the other was estimated using the statistical test defined by Kal *et al*.^3^ All p-values were adjusted for multiple comparisons using Bonferroni corrections. Analysis by operon: for this analysis, we took into account possible polar effects created by the Tn-insertions into genes transcribed from a single promoter and considered that fitness is attributable to the inactivation of operons rather than individual genes.

##### Murine models of bacterial neonatal severe infection

###### Bacterial preparations

The saturated bank of *E. coli* K1 S88 was grown overnight in LB and then diluted in sodium phosphate buffer, pH 7·6, in an appropriate volume according to the needed bacterial concentration for challenge. *E. coli* K1 S88, *E. coli* K1 E11, *E. coli* K1 E11 *Δpga, E. coli* K1CTX-M, Group B *Streptococcus* NEM316 were grown overnight at 37°C on a blood agar plate. From that plate, a 150mL flask containing 15 mL of LB was inoculated to reach an Optical Density (O.D) at 650 nm=0·1 that was incubated at 37°C for about 2 hours, until mid-exponential growth phase (O.D 650 nm=0·5) was obtained. This suspension was administrated directly or was pelleted and resuspended in sodium phosphate buffer pH 7·6, in an appropriate volume according to the bacterial concentrations needed for the challenge.

###### *E. coli* K1 intraperitoneal challenge model

Two or three day-old CD1 mice were separated from their dams and received an IP injection of the infectious bacterial suspension (5×10^6^ CFU of the Tn-mutants bank in 50μL/ mouse). Green food dye was added to visually used the IP location of the bacterial challenge (Figure S2). The neonates were routinely monitored every 2-6 hours and humanely euthanized if they showed signs of severe illness. All surviving animals were euthanized at 24 hours after inoculation. The spleens and the livers were harvested and homogenized, serial dilutions of each organ were made and plated on blood agar plates and MacConkey plates to allow the most precise calculation of bacterial load in each organ.

###### Gavage model of neonatal bacterial meningitis

Two or three day-old CD1 mice were separated from their mothers two hours before challenge by oral gavage to limit interference from milk in the induction of infection. The neonatal mice also received an IP administration of Ranitidine (20 μg/mouse in 50 μL of saline solution) to decrease the potential for bacterial killing by gastric acidity. For intragastric administration of bacterial suspensions (5×10^6^ CFU of the Tn-mutants bank or 10^5^ to 5×10^6^ CFU of a specific bacterial strain in 50μL/ animal) to the neonates, anesthesia using inhaled Isoflurane was performed. Green food dye in the inoculum was used to confirm the intragastric location of the bacterial challenge (Figure S2). Infected neonates were monitored and euthanized as described above, and bacterial burdens in spleens, livers and brains also determined as described above. Litters of ten pups were used both for the IP and the gavage models.

##### Important genes for systemic dissemination and brain infection

At least 3×10^6^ bacteria were recovered from livers of individual mice in both neonatal infection models and from brains of individual mice following oral gavage leading to meningitis. Genomic DNA was extracted, digested by MmeI and gel-sized fragments ligated to oligonucleotides adapters, PCR amplified and sequenced.

##### Construction of the E. coli K1 E11 Δpga deletion mutant

Deletion of the *pga* locus in *E. coli* K1 strain E11 was constructed as previously described.^4^ A kanamycin resistance cassette flanked by FLP recombinase recognition target sites and homology arms to replace the DNA segment of interest were generated by PCR with adequate deletion primers (Table S5). Recombination with the targeted chromosome sequences was mediated by the Red recombinase encoded on pRedET,^5^ resulting in the replacement of the targeted sequence with a kanamycin-resistant cassette; allele replacement was confirmed by PCR. Subsequently, the kanamycin marker was removed, using the FLP expression plasmid pCP20 (Gene **158:**9-14).

##### Antibodies

Two fully human IgG1 MAb with obtained by cloning V-regions from human B cells encoding specificity for either *P. aeruginosa* alginate, MAb F429, or PNAG, MAb F598, were both cloned into the TCAE 6·2 expression vector providing with identical human lambda-light chains and gamma-1 heavy chain constant regions. Both have been described previously.^6,7^

##### Animal sera

Goat antiserum was raised to deacetylated PNAG, as previously described,^8^ using a maleimide-sulfydryl–based conjugation scheme. In brief, purified dPNAG was derivatized with a maleimide group using the chemical linker *N*-[γ-maleimidobutyryloxy] succinimide ester (GMBS). Free sulfhydryl (SH) groups were introduced into the carrier protein TT by treatment with *N*-succinimidyl-3-(2-pyridyldithio)-propionate (SPDP). The maleimide-dPNAG and the SH-TT were mixed together for 2 hours at room temperature to chemically couple these 2 components together, and the dPNAG-TT conjugate was purified by size exclusion chromatography on a Superose 6 column (GE Healthcare®). Goats were immunized as described previously.^8^ This serum was replaced during the course of this study by polyclonal goat antiserum raised to a conjugate vaccine comprised of a nonameric β-1-6-linked glucosamine oligosaccharide conjugated to tetanus toxoid (9GlcNH_2_-TT) that became available after being described in the reference.^9^

##### Confocal microscopy and immunochemical detection of PNAG

###### PNAG expression on different *E. coli* and GBS strains

Microbial samples from broth were fixed by suspension in 4% paraformaldehyde, then spotted onto microscope slides, air-dried, and covered for about one minute with ice-cold methanol. After washing, slides were reacted with control MAb F429 or PNAG-specific MAb F598 directly conjugated to Alexa Fluor 488 at 5·2 μg/mL along with a DNA-visualizing dye (4 μM of Syto 83). After 2 hours at room temperature or overnight at 4 °C, slides were washed and observed by confocal microscopy. For enzymatic treatments to confirm PNAG identification, samples fixed to slides were incubated in Tris-buffered saline (pH 6·4) containing either 50 μg/mL dispersin B (specifically digests the β-(1-6)-*N*-acetyl-glucosamine linkages in PNAG) or 50 μg/mL chitinase (no effect on PNAG antigen, digests the β-(1-4)-*N*-acetyl-glucosamine linkages in chitin) overnight at 37 °C. After washing, cells were treated with the Alexa Fluor 488 directly conjugated to MAb. Samples were washed and mounted for confocal microscopy.^10,11^

###### PNAG expression in brains during bacterial meningitis

20 μL of *E. coli* K1 positive brain homogenate samples were fixed by suspension in 4% paraformaldehyde then spotted onto microscope slides, air-dried, and covered for 1 minute with ice-cold methanol. After washing, slides were reacted with control MAb F429 or PNAG-specific Mab F598 directly conjugated to Alexa Fluor 488 at 5·2μg/mL and 1:200 dilution of rabbit anti-*E. coli* polyclonal followed by 1:250 donkey anti-rabbit AF568. Samples were washed and mounted for confocal.

##### Opsonophagocytosis killing assays

The opsonophagocytic assays followed published protocols.^12,13^ Both a polyclonal goat antiserum raised to a conjugate vaccine comprised of a nonameric β-1-6-linked glucosamine oligosaccharide conjugated to tetanus toxoid (9GlcNH_2_-TT), as well as the fully human IgG1 MAb to PNAG, F598, were used in opsonic killing assays. Normal goat serum (NGS) and MAb F429 to *P. aeruginosa* alginate were used as controls.^7^ Polyclonal sera were first heated at 56°C for 30 minutes. To achieve antigenic specificity in the polyclonal sera, all antisera were adsorbed with ∼10^10^ CFU/mL of *E. coli* K1 Δ*pga* mutant to remove antibodies reactive with antigens on the surface of the bacteria other than those to PNAG. The actual phagocytic killing assay was performed by mixing 100 μL of the PMN suspension, target bacteria, dilutions of test sera or MAb, and a complement source. The reaction mixture was incubated on a rotor rack at 37°C for 90 min. The percentage of killing was calculated by determining the ratio of the number of CFU surviving in the tubes with bacteria, PMN, complement, and test antibodies to the number of CFU surviving in tubes containing control antibodies (MAb F429 or NGS), bacteria, complement, and PMN. Sera were classified as positive if, after subtraction of the background killing in control sera using NGS or an irrelevant MAb, more than 30% of the bacterial CFU were killed, and classified as negative if 30% or less killing was obtained.

##### Passive protection studies

Groups of five to ten newborn mice (CD1 or Complement-C3 deficient) were injected IP with 50 µl of dPNAG-specific goat antiserum to 9GlcNH_2_-TT or NGS, or with 50 µg of human MAb F598 to PNAG or the irrelevant MAb F429, 24 hours before challenge with 10^5^ to 5×10^6^ *E. coli* K1 strains. Twenty-four hours after oral gavage with *E. coli* K1 E2, E11 or S88, mice were sacrificed and the CFU per brain, liver or spleen were determined as described above. In mice passively immunized then challenged with *E. coli* CTX-M, a mortality model up to 24 hours was used to study the antibodies efficacy. This same model has been used to test the protection conferred by antibodies to PNAG in mice challenged with GBS NEM316.

##### *Passive protection following co-infection with* E. coli *K1 and* S. aureus

Two or three day-old CD1 mice were challenge by gavage with different strains of *E. coli* K1 (as previously described). At the same time, an IP administration of 10^5^ to 10^6^ *S. aureus* (either *S. aureus* USA 300 LAC or *S. aureus* PS80) was performed. Three hours after the *S. aureus* IP administration, neonatal mice were injected IP with 50 µl of a dPNAG-specific goat antiserum to 9GlcNH_2_-TT or NGS, or with 50 µg of human MAb F598 to PNAG or irrelevant human MAb, F429. A mortality model at 24 hours following this treatment was used to study the antibodies efficacy. No change in offspring numbers, weight, size were found in pups born from immunized mothers.

##### Maternal immunization and neonatal protection

C3H/HeN female mice (6 to 8 weeks old) were immunized subcutaneously (s.c.) either with dPNAG-TT or Capsular Polysaccharide 5-TT (CP5) given once a week for 3 weeks. Immunization was confirmed by the detection of antibodies to PNAG by ELISA. Neonatal mice were then challenged by gavage with *E. coli* K1 (as previously described). Twenty-four hours after the bacterial challenge, mice were sacrificed and the CFU of *E. coli* present in the blood and brain were determined as described above. In the second part of the experiments, mice born to CP5-TT immunized dams were placed at birth either in cages with a dPNAG-TT immunized dam or a different CP5-TT immunized dam. One week later, infant mice were challenged by oral gavage with *E. coli* K1. Twenty-four hours after the bacterial challenge, mice were sacrificed and the CFU of *E. coli* present in the blood and in the brain were counted.

##### ELISA detection of antibodies to PNAG

ELISA experiments were performed as previously described.^8^ Briefly, ELISA plates (Maxisorp, Nalge Nunc International, Rochester, NY, USA) were coated with 0·6 μg/ml of purified PNAG diluted in sensitization buffer (0·04M PO_4_, pH 7·2) overnight at 4°C.^14,15^ The plates were rinsed three times with PBS, blocked with 5% skim milk for 1 h at 37°C, and rinsed again three times with PBS. Serum samples were then diluted twofold in 5% skim milk with 0·05% Tween 20, incubated for 1 h at 37°C, washed again, and incubated with appropriate alkaline phosphatase-conjugated secondary antibody (Sigma) for 1 h at 37°C. After washes, plates were incubated with *p*-nitrophenyl phosphate substrate, and the optical density at 405 nm (OD_405_) was determined by an enzyme-linked immunosorbent assay (ELISA) plate reader (BioTek Instruments, Winoski, Ill.) after a 1-h incubation.

##### E. coli *interactions with brain microvascular endothelial cells*

###### Cell cultures

Two different cell lines of human brain microvascular endothelial cells were used in this study. Human brain micro-endothelial cells (HBMEC), immortalized by SV-40 large T antigen transformation, and maintaining the morphological and functional characteristics of the primary endothelium, were obtained from K.S. Kim, MD (Johns Hopkins University, Baltimore).^16^ HBMEC were grown on collagen-coated tissue culture plates in RPMI medium (Sigma-Aldrich) containing 10% heat inactivated fetal bovine serum, 10% Nu-Serum, 2 mM glutamine, 1 mM pyruvate, 100 U/mL penicillin, 100 μg/mL streptomycin, 1% MEM nonessential amino acids and vitamins at 37 °C in humid atmosphere of 5% CO2 until confluence (about five days). The Human Cerebral Microvascular Endothelial Cell line (hCMEC/D3) (provided by Cedarlane, and developed by P.O Couraud, INSERM, Paris) was prepared from cerebral microvessel endothelial cells by transduction with lentiviral vectors carrying the SV40 T antigen and human telomerase reverse transcriptase.^17^ hCMEC/D3 were grown on collagen type I coated T75 flasks in EndoGRO-MV Culture MediumTM (provided by EMD Millipore) containing 5% heat inactivated fetal bovine serum, 0·2% EndoGRO-LS Supplement, 5ng/mL of recombinant human Epidermal Growth Factor, 10 mM of L-glutamine, 1μg/mL of hydrocortisone hemisuccinate, 0·75 U/mL of heparin sulfate, 50 μg/mL of ascorbic acid, 200 ng/mL of human basic Fibroblast Growth Factor, 100 U/mL penicillin, 100 μg/mL streptomycin, at 37°C in humid atmosphere of 5% CO2 until confluence (about five days). As for the HBMEC, cells reaching confluence were harvested (using trypsin) and either transferred to a 24-wells plate for further experiments or cryopreserved at -150°C.

###### Invasion, binding and translocation assays

Confluent HBMEC or hCMEC/D3 monolayers grown in 24-well plates were infected with bacteria at a multiplicity of infection (MOI) of 100 alone, or with addition of either (i) goat antibody to dPNAG or NGS (diluted 1/10 or 1/100); (ii) with MAb F598 or MAb F429 (100 μL of a 1 mg/mL solution). After 90 min of incubation at 37°C, the monolayers were washed with the appropriate medium depending on the cell line and then incubated with gentamicin (100 μg/ml) for 1 hour to kill extracellular bacteria and washed again. Finally, they were lysed by 0·1% Triton-X100 in PBS and the released intracellular bacteria enumerated. The invasion frequency was calculated by dividing the number of internalized bacteria by the original inoculum. To measure binding, the assay was performed as above except that the gentamicin treatment step and the cell lysis by Triton-X100 were omitted and monolayers were washed at least 8 times by PBS to insure removal of unbound bacteria. For translocation assays, HBMEC or hCMEC/D3 monolayers were established on transwells, then infected with bacteria at a MOI of 100 alone, or with antibodies and translocation levels measured at 0 and 30 minutes.

###### Trans-endothelial electrical resistance

HBMEC were cultured to confluence then TEER was measured using a Millicell-ERS (Millipore, Burlington, MA, USA). The collagen-treated transwell inserts were used to measure the background resistance. In the control monolayer no bacteria was added. Values were expressed as *ω* × cm^2^ and were calculated by the formula: [the average resistance of experimental wells − the average resistance of blank wells] × 0·33 (the area of the transwell membrane). The TEER values were recorded in real-time every 30 minutes in triplicates for 90 minutes.

##### GBS experiments

###### Attempts to truncate gene GBS1605 by double crossing over

Deletion of the *pgaC*-like (gene GBS1605) was attempted in *S. agalactiae* NEM316 by allelic exchange with a truncated copy of GBS1605 cloned into the temperature-sensitive shuttle vector, pJRS233, following a procedure previously described.^18^ Approximately 900 bp fragments upstream and downstream of GBS1605 were amplified by PCR using NEM316 chromosome as the template and primer pairs GBSpgaC1/GBSpgaC2 and GBSpgaC3/GBSpgaC4. The forward primer binding to the 3’-end of *pgaC* and the reverse primer to the 5’-end were modified to carry a *BamH1* restriction site while GBSpgaC1 and GBSpgaC4 primers were modified to carry a *pst1* and *xho1* site, respectively. Following restriction, ligation and re-amplification using GBSpgaC1 and GBSpgaC4, the resulting fragment carrying the truncated GBS1605 gene (deletion of ca. 50% of the mid-section of the allele) was cloned in *xho1/pst1* double digested temperature-sensitive shuttle vector pJRS233 to create plasmid pJRS233pgaC-KO (Table S4). After electroporation and amplification in *E. coli* DH5-alpha, the recombinant plasmid was recovered (Qiagen plasmid extraction MIDI) and introduced into the chromosome of NEM313 by electro-transformation and homologous recombination followed by excision as described.^18^ Multiple attempts (confirmation PCR on 1000+ colonies using *pgaC* flanking primers) yielded the same negative result: the conservation of a wild-type GBS1605 allele in the chromosome and the excision and loss of the truncated allele and the pJRS233 backbone.

###### Nisin inducible antisense RNA

For antisense expression, we cloned a 721 bp complementary 3’ end fragment of GBS1605 (primers pgaC-AS1 and pgaC-AS2), including the native transcription start site but avoiding any overlap between adjacent genes, in the antisense orientation downstream of the inducible promoter (PnisA) of pMSP3535.^19^ Following restriction digestion, the GBS1605 amplicon was cloned into pMSP3535 for expression in the antisense direction under nisin induction. The ligated construct was initially transformed into *E. coli* DH5-alpha and sequence-verified. Purified recombinant plasmid was then transformed into *S. agalactiae* NEM316 followed by plating on Todd Hewitt agar supplemented with 10 μg/mL erythromycin. Growth curves were carried out utilizing NEM316 strains with gene-specific antisense fragment or empty vector as follows: Overnight cultures were diluted 1:100 in Todd Hewitt broth supplemented with 5 μg/mL erythromycin. At an approximate O.D. 600 of 0·2, cultures were diluted such that approximately 100 cells were inoculated in triplicate into inducing (0·1 to 5 μg/mL) or non-inducing Todd Hewitt broth in a microtiter plate. Growth was followed for 24 h by measuring O.D.600 on a Biochrom WPA spectrophotometer (Biochrom, Holliston, MA).

For qRT-PCR, total *S. agalactiae* NEM316 RNA was isolated from exponentially growing cells, with or without nisin induction in Todd Hewitt broth, using the RNeasy midi kit (Qiagen) as recommended by the manufacturer. cDNA was synthesized from total RNA (∼1·5 mg) by using the Superscript III First-Strand Synthesis System (Invitrogen) according to the manufacturer’s instructions. Using synthesized cDNAs, qRT-PCR was performed using Express SYBR GreenER qPCR Supermix (ThermoFisher, Waltham, MA, USA) and a real-time PCR cycler system (Rotor-Gene Q5-Plex; Qiagen, Hilden, Germany) with the following program: 95 C for 10 min, and subsequently 40 cycles of 95 C for 15 sec, 55 C for 1 min. Relative transcript levels were calculated using REST 2009 Software (Qiagen). Expression of elongation factor *tuf* was used as a housekeeping control gene.

##### Statistical analysis

Nonparametric data were evaluated by the Mann Whitney U test for 2-group comparisons and multi-group comparisons by the Kruskal Wallis test with Dunn’s multiple comparison test for pairwise comparisons. Parametric data were analyzed by t-tests (for 2-group comparisons) or ANOVA with Tukey’s multiple comparison test for pair-wise comparisons. Survival analysis utilized the Kaplan-Meier method. All analyses, apart from the Tnseq analyses, used GraphPad Prism 6·0 (GraphPad Software, San Diego, CA).

## Supplemental figures

**Figure S1.**
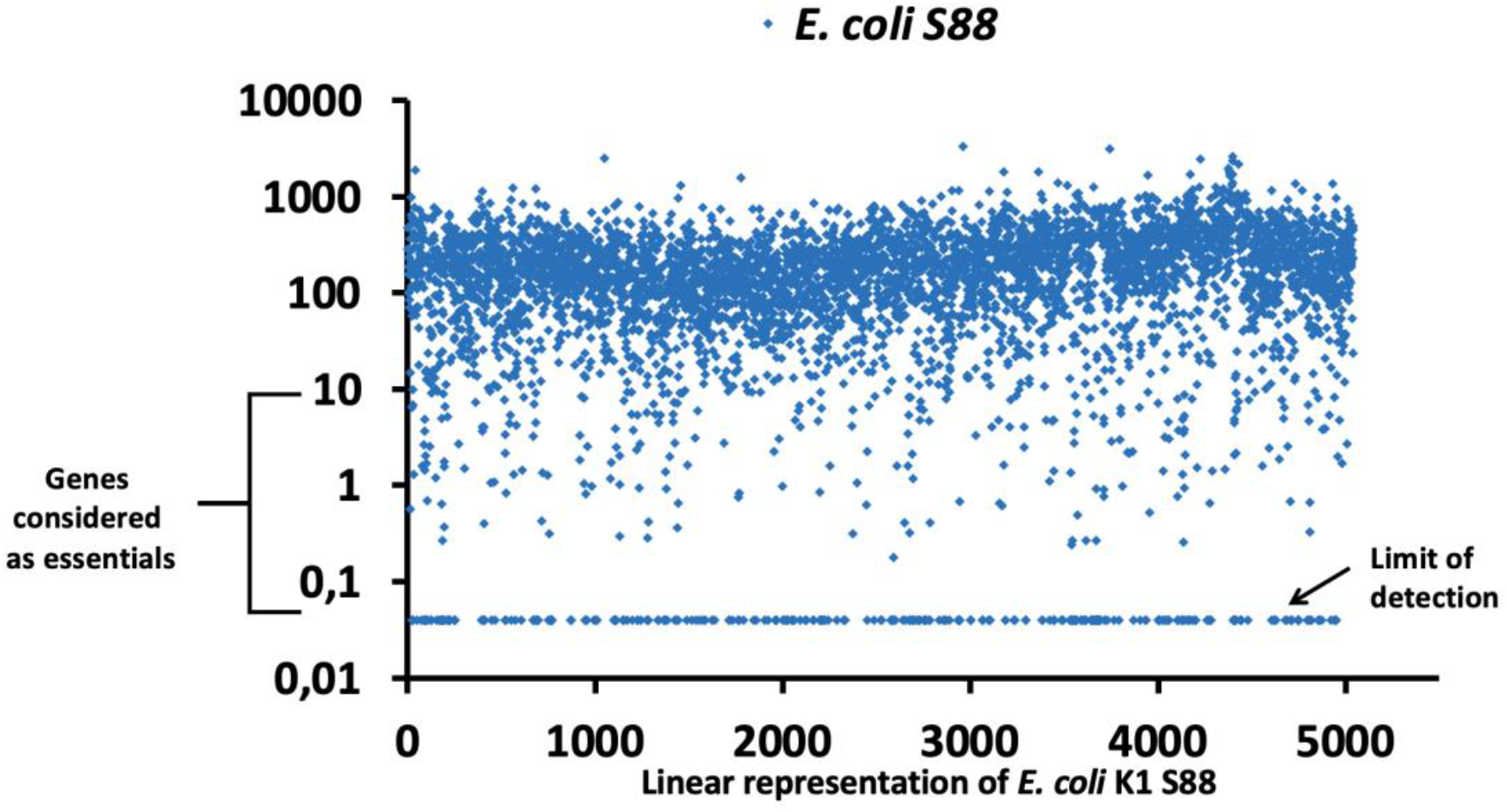
Properties of the *E. coli* K1 S88 TnSeq bank grown in lysogeny broth. Analysis of the overall frequency of Tn insertions into the chromosome was based on use of one million sequencing reads to normalize the data from different DNA preparations (RPKM). In lysogeny broth (LB), the 300·000 *E. coli* K1 S88 Tn-mutants had a relatively homogeneous distribution of Tn insertions across *E. coli* K1 S88 chromosomes with no genes having Tn insertions with more than 10·000 sequencing reads. Following definitions we have previously used, genes with <10 sequencing reads were defined as those essential for growth in LB.

**Figure S2.**
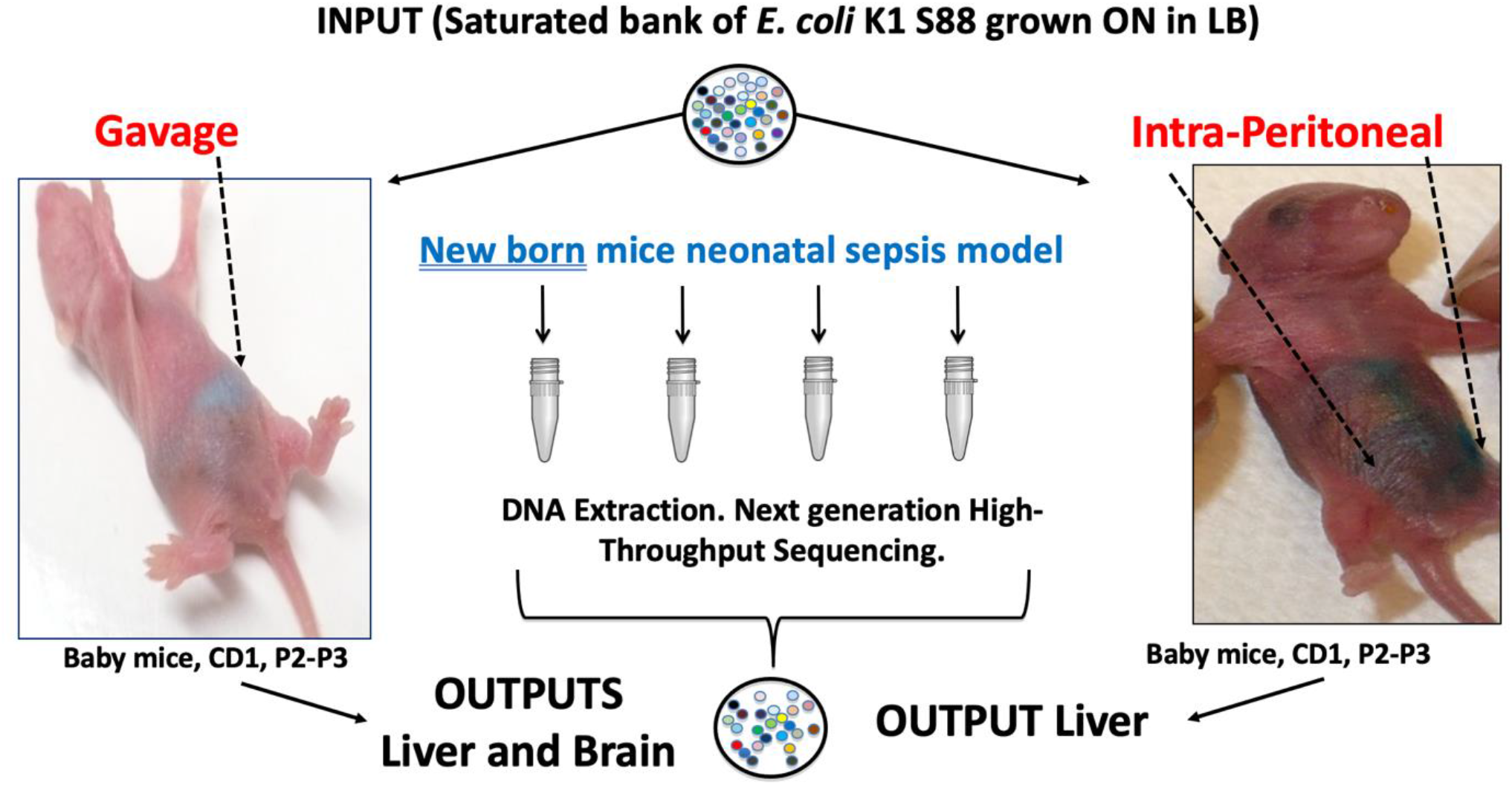
*E. coli* K1, neonatal sepsis model. The 3×10^5^ saturated bank of *E. coli* K1 S88 grown overnight (ON) in lysogeny broth (LB) was administered by an IP injection or by gavage (5×10^6^/mouse). Green food dye was used to confirm the intraperitoneal or intragastric location of the bacterial challenge following IP injection or oral gavage of the inoculum. To determine the systemic dissemination in both routes of infection, newborns’ livers were harvested 24 hours after the challenge. To study brain infection in the gavage model of neonatal bacterial meningitis, brains were harvested 24 hours after the bacterial challenge. At least 3× 10^6^ were recovered from each organ of individual mice. Genomic DNA was extracted, digested by MmeI and gel-sized fragments ligated to oligonucleotides adapters, PCR amplified and sequenced.

**Figure S3.**
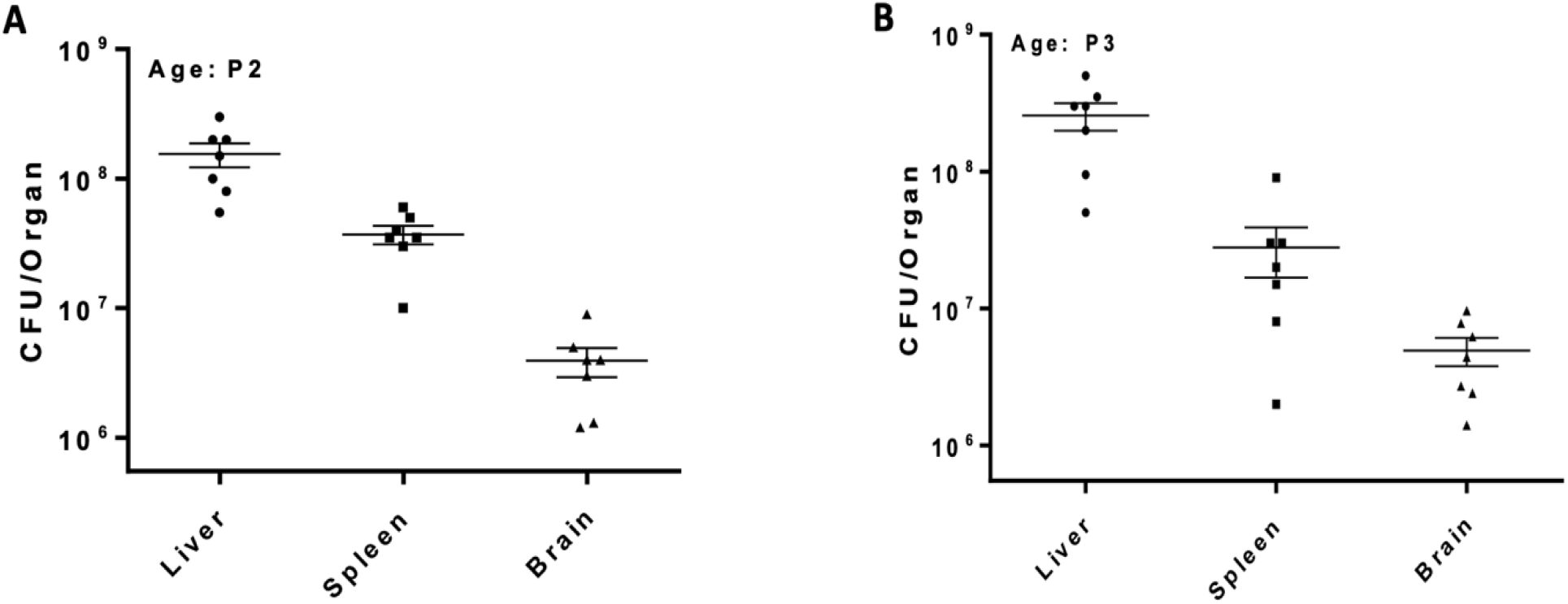
Systemic dissemination of *E. coli* K1 after intraperitoneal infection of new born mice. Strain: *E. coli* K1 S88. Inoculum: 4×10^6^ Age of the new born mice: 2 days (A) and 3 days (B) old Bars represent the mean, and error bars depict the 95% confidence interval.

**Figure S4.**
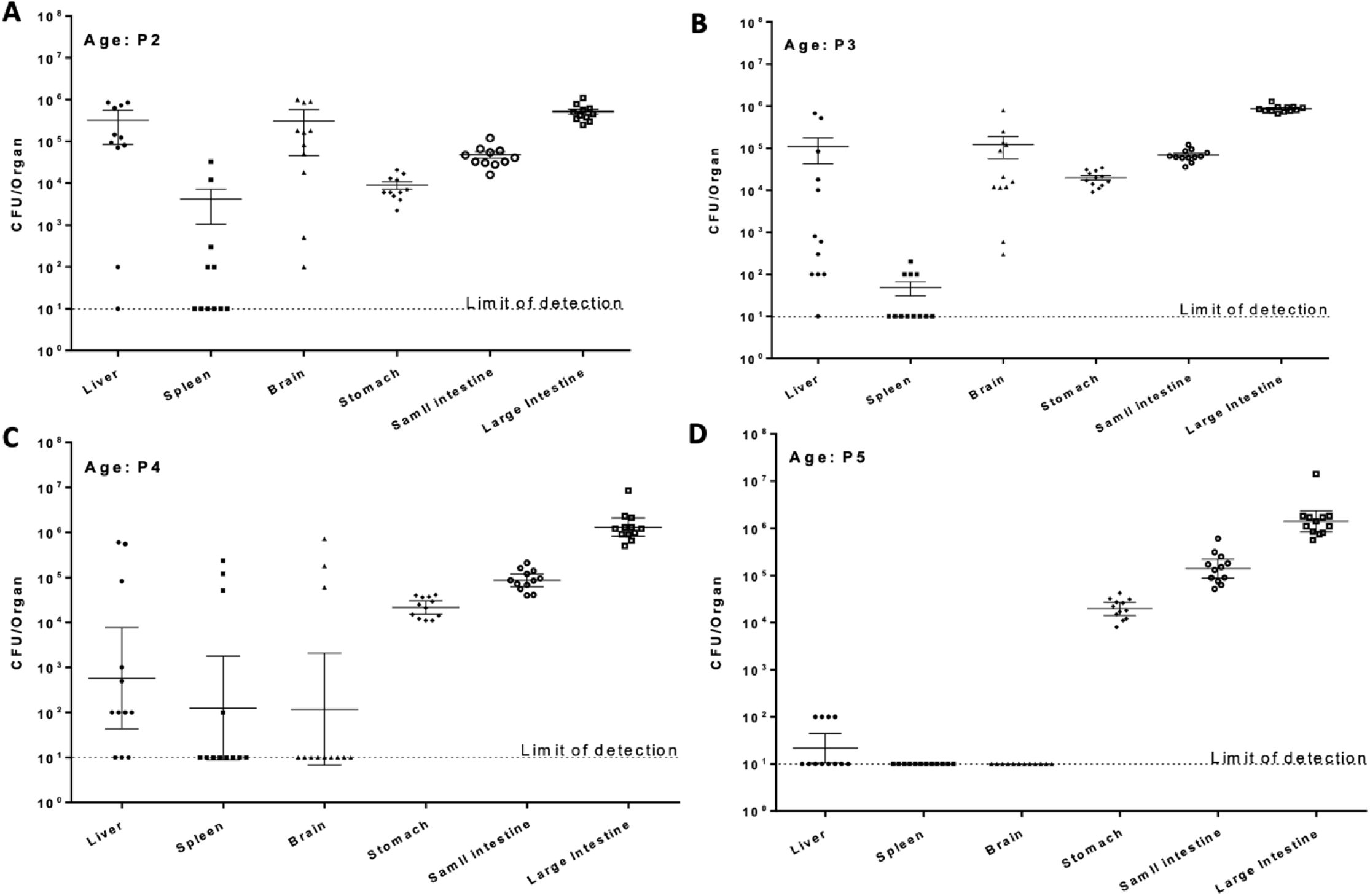
Systemic dissemination of *E. coli* K1 from the gastro-intestinal tract after gavage of new born mice. Strain: *E. coli* K1 S88. Inoculum: 2×10^6^ Age of the neonates: 2 days (A), 3 days (B), 4 days (C) et 5 days old (D) Bars represent the mean, and error bars depict the 95% confidence interval.

**Figure S5.**
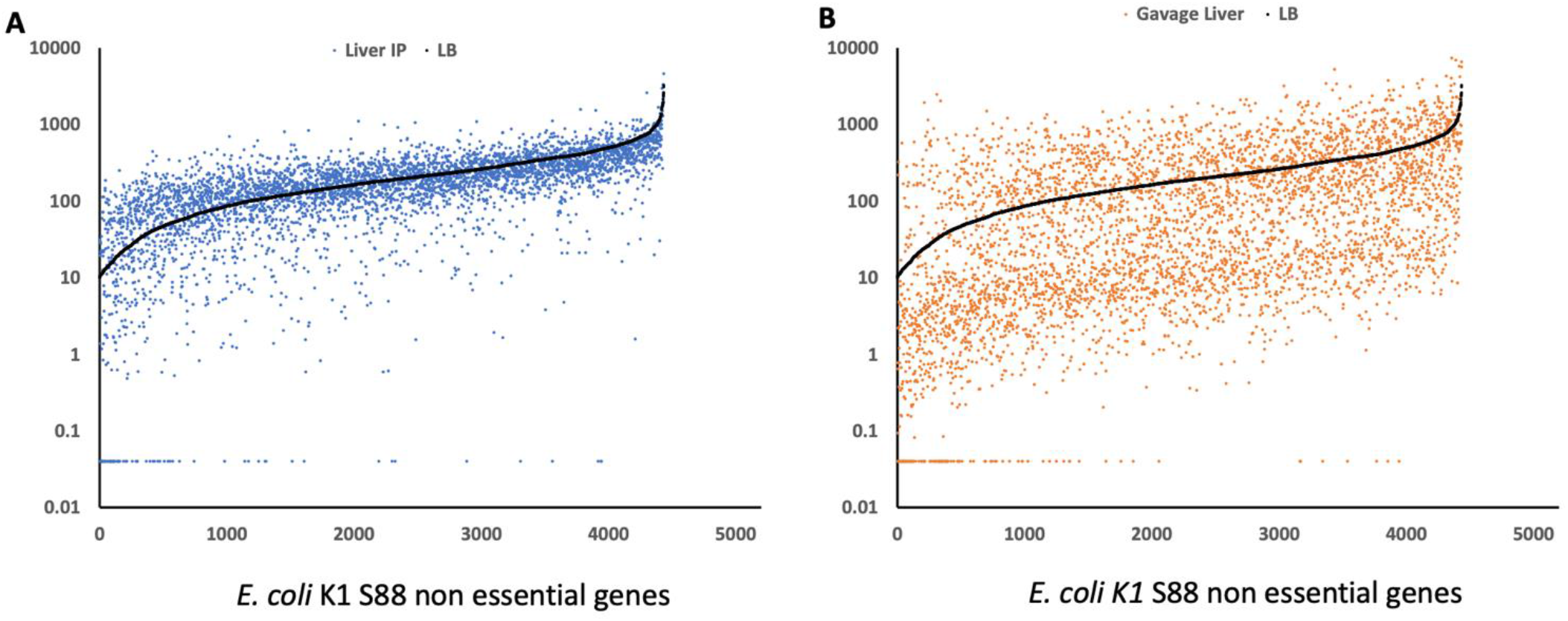
*In vivo* fitness of *E. coli* K1 S88 Tn library. Relative ranking and absolute number of RPKM that changed for 4818 non-essential genes of *E. coli* K1 S88 in the new born mice meningitis models, comparing the RPKM in the lysogeny broth (LB) input with those obtained in the liver after IP challenge (A) or gavage (B). Dots above the input lines (in black) indicate Tn insertions in genes with a positive fitness (increased RPKM), whereas dots below the input line indicate those with a negative fitness (decreased RPKM).

**Figure S6.**
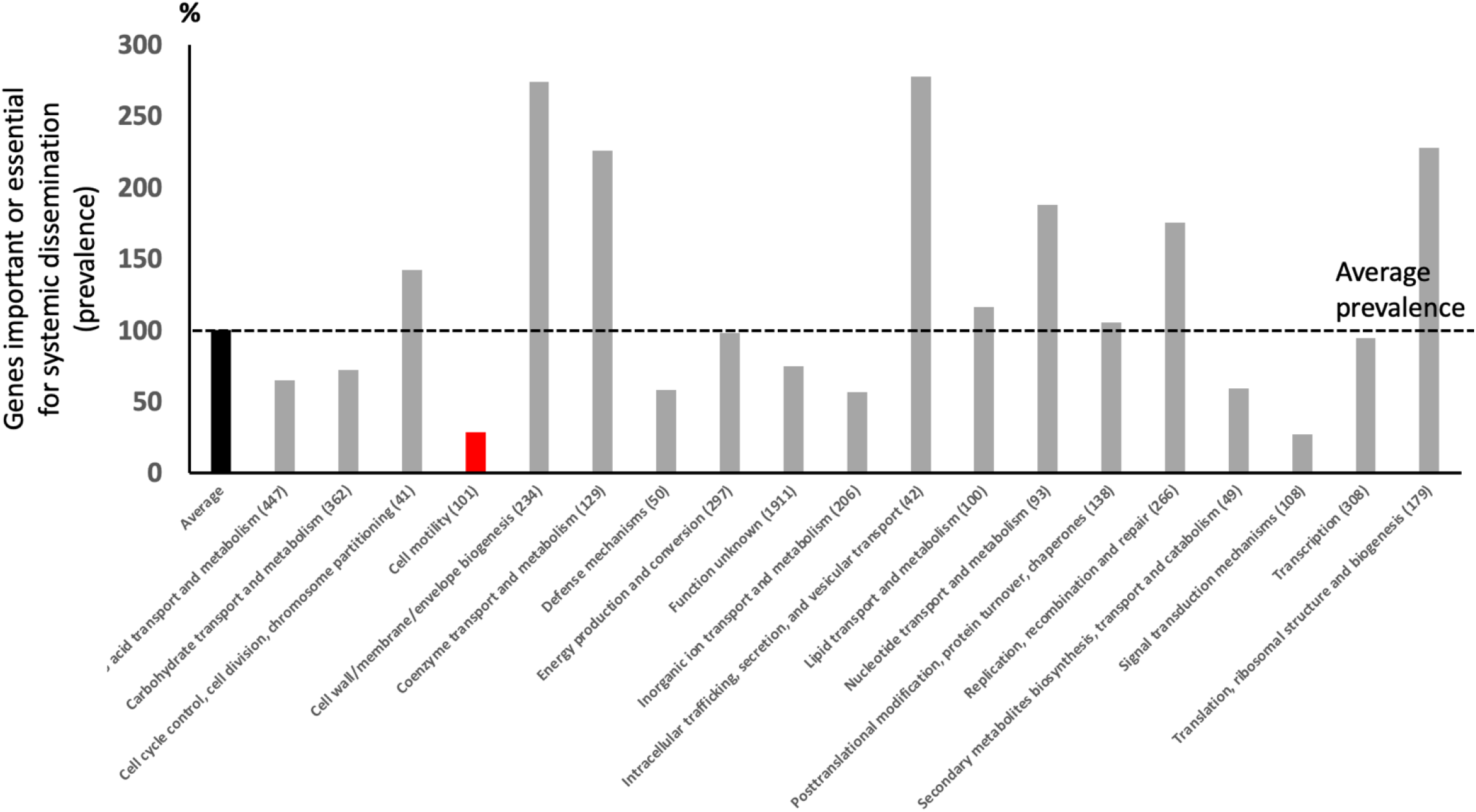
*E. coli K1* important genes for systemic dissemination analyzed by functional classes. *E. coli* K1 S88 genes were classified into 19 functional classes (https://www.genome.jp/kegg/). The prevalence of genes important or essential for systemic dissemination (defined by more than 10 fold decrease in both routes of infection between the growth in lysogeny broth and the mutants in these genes recovered from the livers) in each functional class is represented, compared to the overall average prevalence of these genes in *E. coli* K1 S88. The red bar represent the importance of the genes belonging to the functional class “motility” in the systemic dissemination.

**Figure S7.**
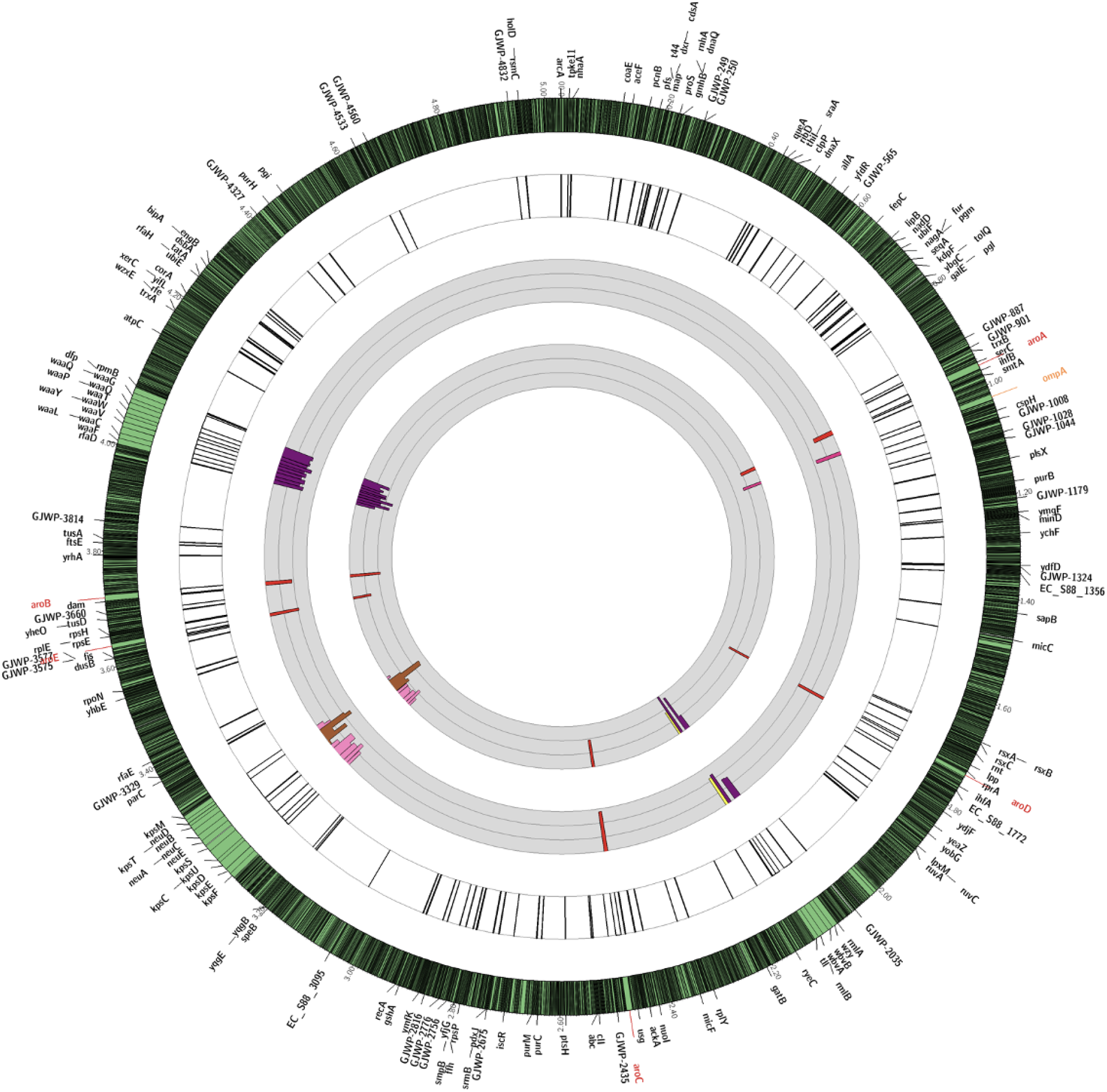
Important genes for systemic dissemination: localization on *E. coli* K1 S88 chromosome and quantitative analysis. The outermost circle represents the full *E. coli* K1 S88 genome with a 30-times magnification of the regions of interest. The first gray circle represents the fold change of the reads of the Tn insertions from the LB to the Liver after IP challenge. The second gray circle represents the fold change of the reads of the Tn insertions from the LB to the Liver after gavage. The thin grey circular lines represent 10-fold-changes (i.e., at log_10_ scale). The limit of the fold change representation is 1000. Bars represent changes in individual genes. Bars pointing toward the circle’s center represent Tn interrupted genes resulting in negative fitness. Major, previously described, virulence factors of *E. coli* K1 are highlighted in color: LPS and K1 capsule encoding genes as examples for genes clusters, OmpA as example for a unique gene and Aro encoding genes as example of several individual genes (i.e not in cluster) located on several and distanced part of the chromosome.

**Figure S8.**
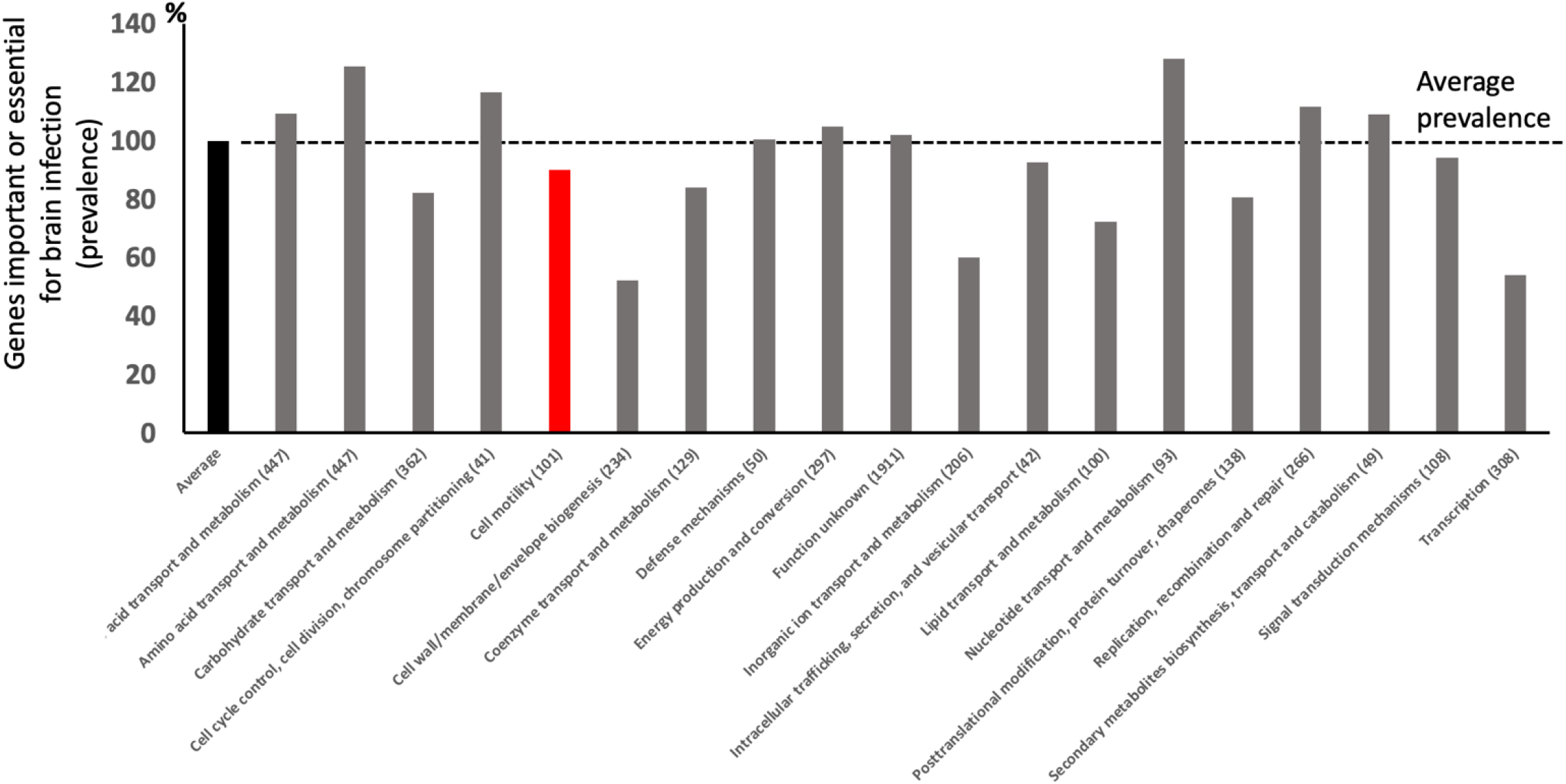
Genes important for brain infection, analysis by functional classes. *E. coli* K1 S88 genes were classified into 19 functional classes (https://www.genome.jp/kegg/). The red bar represent the importance of the genes belonging to the functional class “motility” in the brain infection. No difference between this functional class and the other functional classes is found.

**Figure S9.**
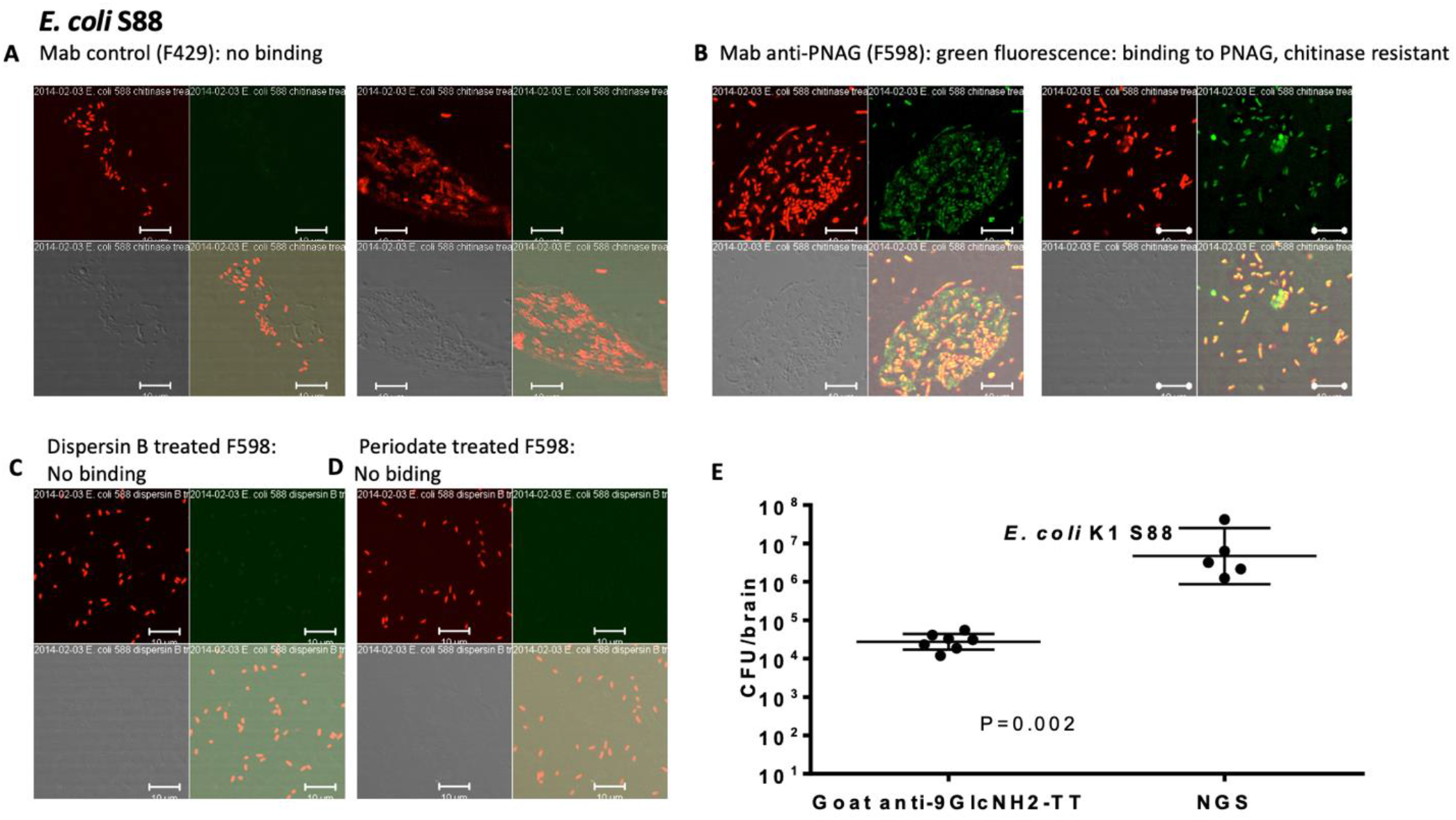
*E. coli* K1 S88 and PNAG. (A-D) Detection of PNAG production (green) by *E. coli* K1 using confocal microscopy to visualize binding of MAb F598 to PNAG. Control: MAb, F429, to *P. aeruginosa* alginate. The chemical properties of the PNAG antigen were confirmed using a previously reported approach by enzymatic digestion with Dispersin B (sensitive) and Chitinase (41). If PNAG is present, green fluorescence is dramatically decreased after Dispersin B treatments. Chemical specificity was confirmed using periodate (sensitive). (E) Prophylactic (24 h pre-challenge) effect of 50 µl of opsonic goat polyclonal antibodies to the synthetic oligosaccharide 9GlcNH2-TT on *E. coli* K1 levels in the brain 24 h after challenge by gavage. Controls received normal goat serum (NGS). Bars represent the mean, and error bars depict the 95% confidence interval. P values determined by nonparametric t-test.

**Figure S10.**
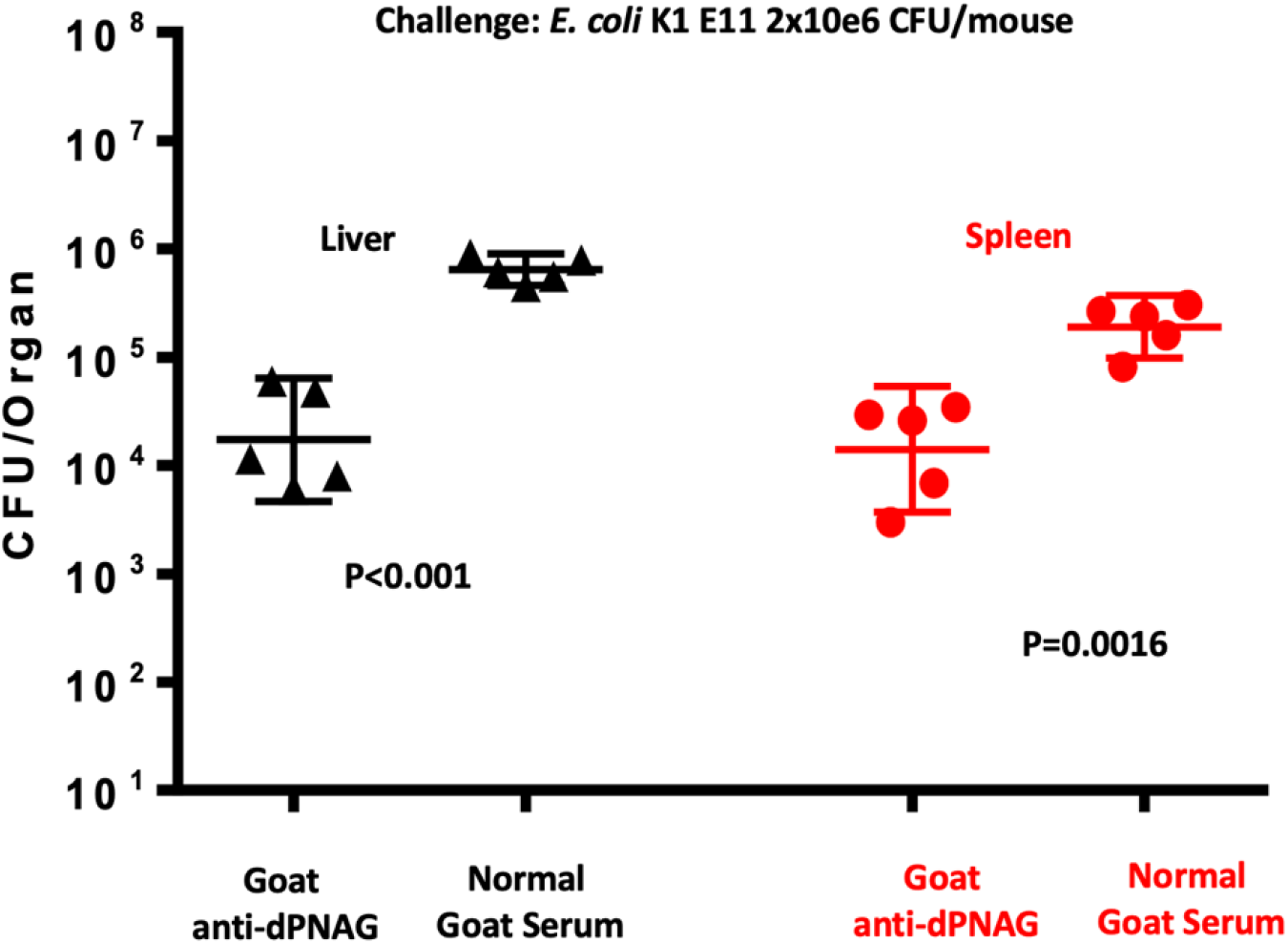
Prophylactic effect of polyclonal antibodies to PNAG on *E. coli* K1 levels in liver and spleen. Prophylactic (24 h pre-challenge) effect of 50 µl of opsonic goat polyclonal antibodies to PNAG on *E. coli* K1 levels in the liver and the spleen 24 h after challenge by gavage. Controls received normal goat serum (NGS). P values determined by nonparametric t-test. Bars represent the mean, and error bars depict the 95% confidence interval.

**Figure S11.**
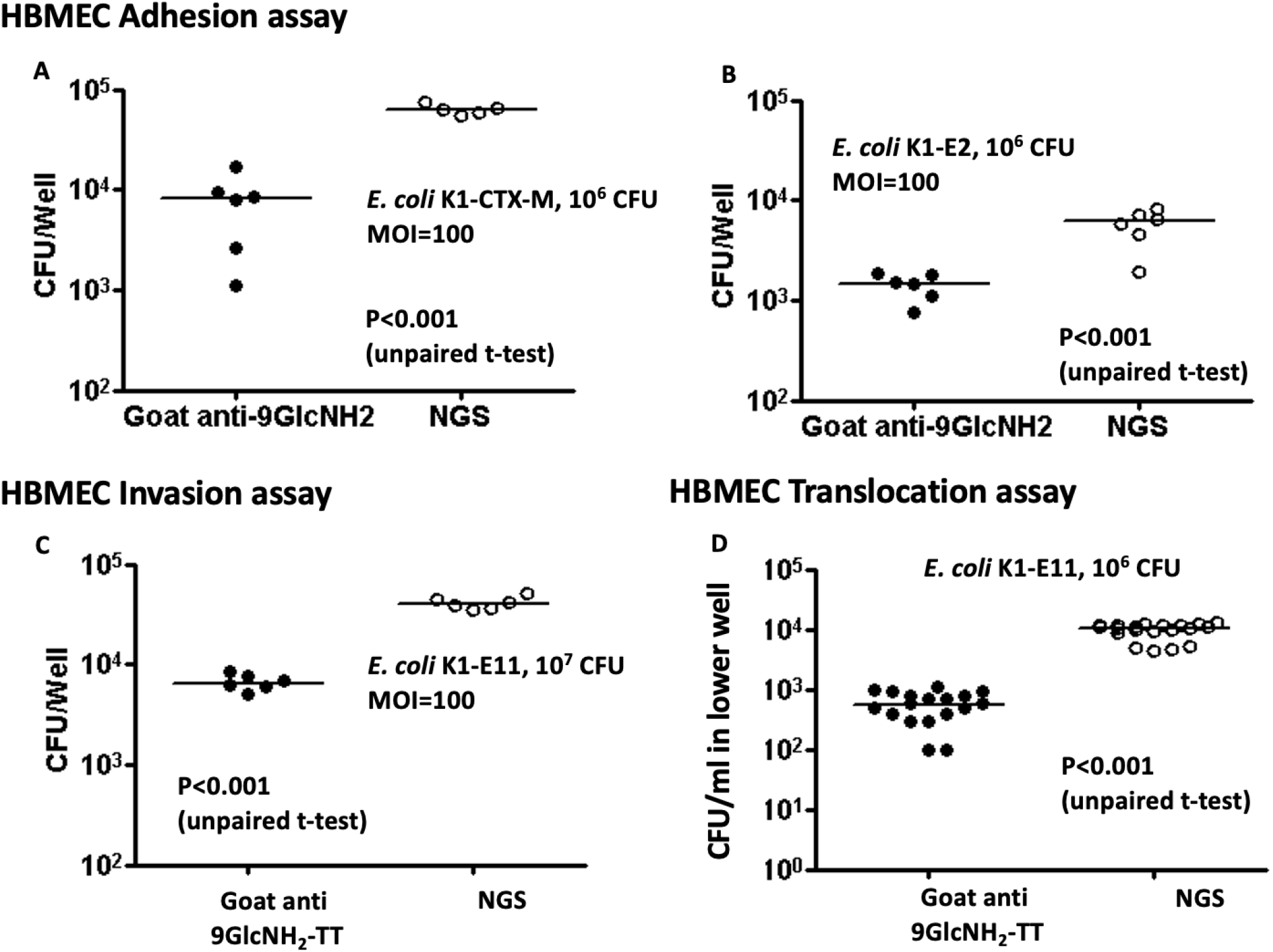
Adhesion (other strains), Invasion and translocation assays. Goat polyclonal antibodies to the synthetic oligosaccharide 9GlcNH2-TT significantly inhibits adherence, invasion and translocation of different strains of *E. coli* K1 applied to HBMEC compared to normal goat serum (NGS). (A) Adhesion assay of *E. coli* K1-CTX-M. (B) Adhesion assay of *E. coli* K1-E2. (C) Invasion assay of *E. coli* K1-E11. (D) Translocation assay of *E. coli* K1-E11. Symbols are individual wells, lines medians, P values unpaired t-tests.

**Fig. S12.**
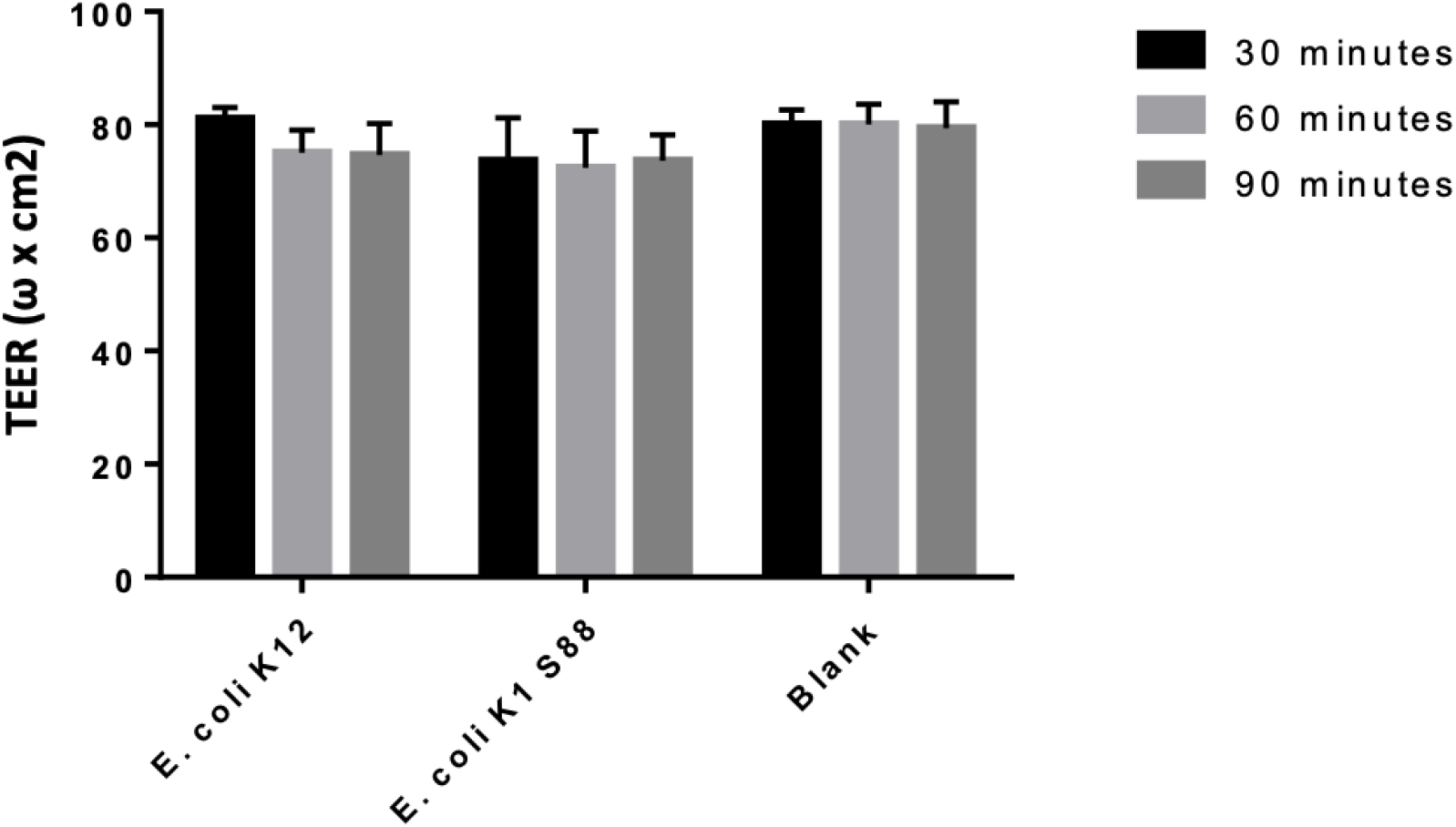
Transendothelial electrical resistance of HBMEC in presence of different strains of *E. coli*. Transendothelial electrical resistance of HBMEC after 30, 60 and 90 minutes of incubation with *E. coli* K12, *E. coli* K1 S88 or blank. Bars represent the mean, and error bars depict the confidence interval

**Figure S13.**
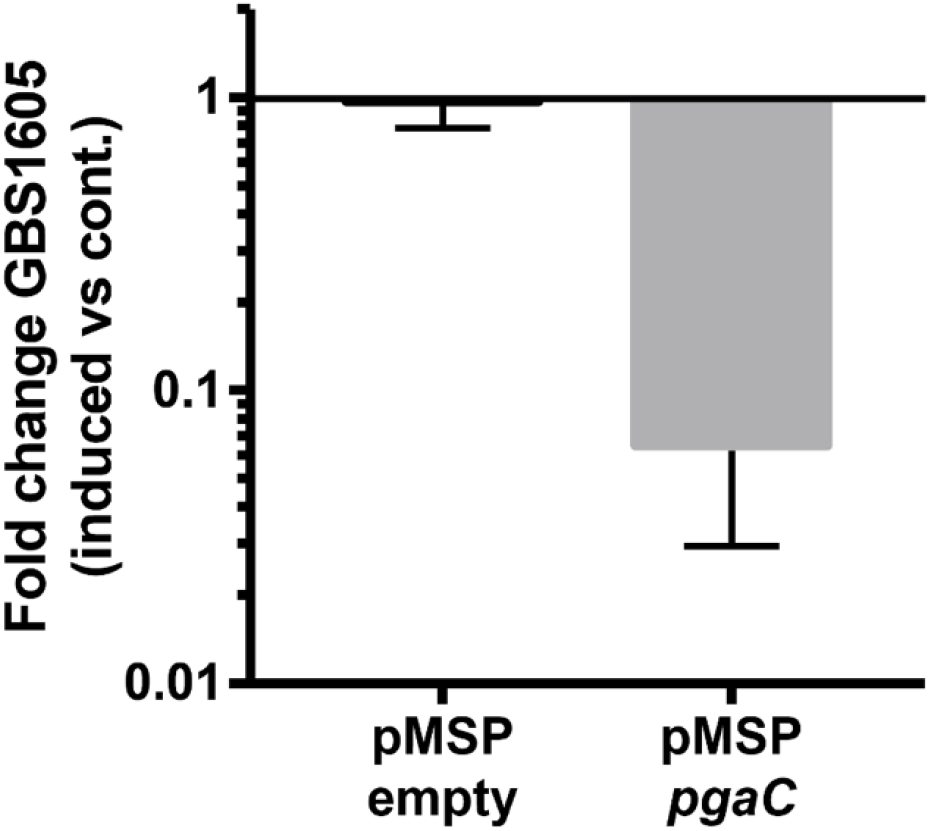
Decrease of GBS1605 transcripts in presence of nisin and asRNA pMSP*pgaC*. qRT-PCR reveals a 30-fold decrease in GBS1605 transcripts in the presence of 0·5 mg/mL nisin induction and the antisense construct pMSP*pgaC*. As a control, presence of empty vector pMSPempty had no significant effect on transcription levels of GBS1605 with or without nisin induction. Transcription changes are expressed in ratio induced versus non-induced. In all tested conditions, GBS1605 relative transcription levels were normalized using those of housekeeping gene tuf. Experiments were performed in three independent biological replicates. Bars represent the mean, and error bars depict the 95% confidence interval.

**Table S1. *E. coli* essential genes for growth in LB**

File Table S1.xslx

**Table S2. Genes important or essential for systemic dissemination**

File Table S2.xslx

**Table S3. Data for Circos Brain 1203 and only outer membrane**

File Table S3.xslx

**Table S5.** Strains used in the study.

| Strain | Source |
| --- | --- |
| *E. coli* K1 S88 | Provided by S. Bonacorsi<sup>20</sup> |
| *E. coli* K12 MG1655 |  |
| *E. coli* LF82 | Provided by H. Sokol<sup>21</sup> |
| *E. coli* K1 E11 | Clinical strain from the Brigham and Women's Hospital |
| *E. coli* K1 E11 $\Delta pga$ | This study |
| *E. coli* E2 | Clinical strain from the Brigham and Women's Hospital |
| *E. coli* K1 CTX-M | Provided by R. Bonnet<sup>22</sup> |
| *S. aureus* PS80 | Provided by J.C Lee |
| *S. aureus* USA300 LAC | <sup>13</sup> |

**Table S6.**
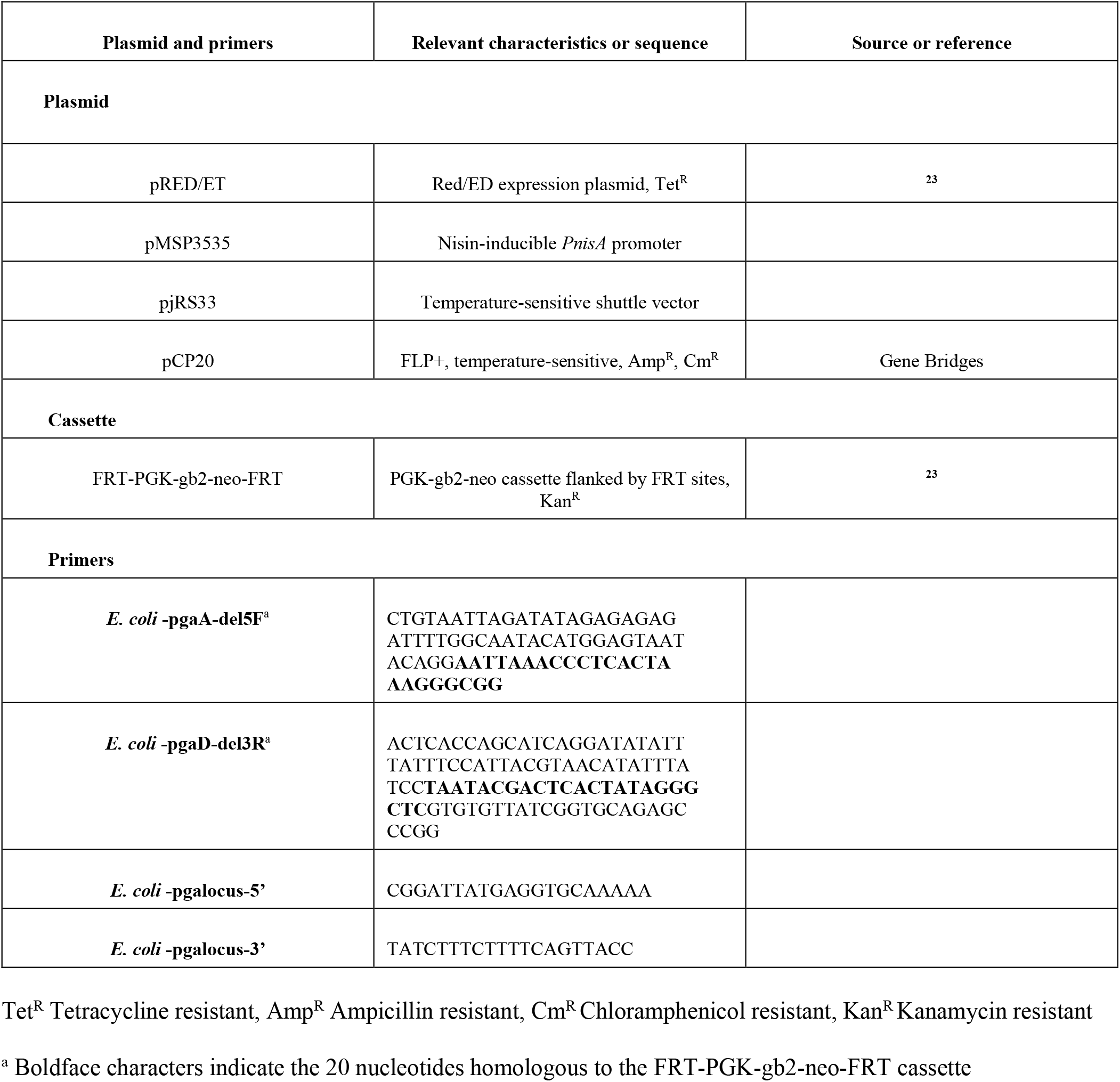
Plasmids and primers used in the study.

